# Specificities of exosome versus small ectosome secretion revealed by live intracellular tracking and synchronized extracellular vesicle release of CD9 and CD63

**DOI:** 10.1101/2020.10.27.323766

**Authors:** Mathilde Mathieu, Nathalie Névo, Mabel Jouve, José Ignacio Valenzuela, Mathieu Maurin, Frederik Verweij, Roberta Palmulli, Danielle Lankar, Florent Dingli, Damarys Loew, Eric Rubinstein, Gaёlle Boncompain, Franck Perez, Clotilde Théry

## Abstract

Despite their important and multiple roles in intercellular communications, the different populations of extracellular vesicles (EVs) and their secretion mechanisms are not fully characterized yet. In particular, how and to what extent EVs form either as intraluminal vesicles of endocytic compartments (exosomes), or at the plasma membrane (ectosomes) remains unclear. We followed in HeLa cells the intracellular trafficking of the EV markers CD9 and CD63 from the endoplasmic reticulum to their residency compartment and identified transient co-localization both at the plasma membrane (PM) and in endosomes, before they finally segregate. CD9 was more abundantly released in EVs than CD63. However, when forcing expression of CD63 at the PM, by mutating its lysosome-addressing motive, its secretion in EVs was increased. Thus, in HeLa cells, small ectosomes are more prominently released than exosomes. By comparative proteomic analysis, we identified a few surface proteins likely specific of either exosomes (e.g. LAMP1) or ectosomes (e.g. BSG, SLC3A2), based on their known intracellular location in lysosomes or the PM, and on the different effects on their release of Bafilomycin A1, a drug that neutralizes endosomal pH. Our work sets the path for molecular and functional discrimination of exosomes and small ectosomes in any cell type.

## INTRODUCTION

All cells release membrane-enclosed vesicles, collectively called Extracellular Vesicles (EVs), in their environment. These EVs contain a selected set of lipids, nucleic acids and proteins from their cell of origin, and thus can transfer a complex array of information to surrounding or distant cells^1,2^. EVs can form by direct outward budding from the plasma membrane (PM) of prokaryotic and eukaryotic cells. In eukaryotic cells, EVs can also form first as intraluminal vesicles of internal multivesicular compartments of the endocytic pathway (MVBs), and are then secreted upon fusion of these compartments with the PM. To clarify the nomenclature, it has been recently recommended to use the term “exosomes” specifically for the MVB-derived EVs, rather than for all small EVs^3^. The PM-derived EVs, on the other hand, are called various names, such as microvesicles, microparticles, or ectosomes: we will use here the latter term which is exclusively used for PM-derived EVs, whereas the others are also used for any type of EVs.

Since they form at different subcellular sites, exosomes and ectosomes will likely contain different sets of specific cargoes, and thus different functions. However, this hypothesis has not been conclusively confirmed so far, given the difficulty to separate exosomes from ectosomes of the same size present in biofluids or cells’ conditioned medium, the lack of specific protein markers to distinguish exosomes from ectosomes, and of molecular machineries and tools with demonstrated full specificity for one or the other^4^.

Several tetraspanins, especially CD63^5^, CD81^5^ and CD9^6^ have been used as markers of exosomes for the last two decades, due to their accumulation in small EVs as compared to whole cell lysates, and to the steady-state accumulation of CD63 in MVBs. More recently, however, their presence in other EVs has been observed. By capture of EVs bearing specifically either CD63 or CD9 or CD81, followed by analysis of their protein composition and enrichment in endosomal markers, we proposed that EVs bearing only CD9 or CD81 but not CD63 probably did not form in endosomes (and were thus ectosomes), whereas those bearing CD63 together with one or the two other tetraspanins may correspond to endosome-derived exosomes^7^. This observation was made using EVs released by primary human immune dendritic cells, which complicated a direct validation of the model by carrying out cell biology analyses.

The current study was initiated to determine the actual exosomal or ectosomal nature of EVs containing the different tetraspanins. We used here HeLa cells, a cellular model amenable to experimental manipulations necessary to address cell biology questions. Our reasoning was that, to identify the subcellular origin of the EVs released by these cells, we needed to follow in a time-controlled manner their intracellular trafficking from their initial synthesis in the endoplasmic reticulum (ER) until their secretion in EVs. We adapted to CD63 and CD9 the Retention Using Selective Hook (RUSH) system, which has been used to follow and control trafficking of numerous transmembrane and secreted proteins, and thus identify novel atypical pathways of secretion^8^. This allowed us also to synchronize the release of CD9- or CD63-containing EVs, in order to determine the composition of a more homogenous mixture of newly synthesized EVs. Our results demonstrate that both CD63 and CD9 can be released in small ectosomes formed at the plasma membrane, and that in HeLa cells, exosomes represent a minor subpopulation of small EVs (sEVs) that bear CD63 together with other late endosomal molecules such as LAMP1/2. Their secretion is specifically susceptible to neutralization of the endosomal pH. Secretion of ectosomes, for which we identified novel specific markers in HeLa cells as BSG and SLC3A2, is conversely insensitive to endosomal pH neutralization. Interestingly, CD81, another tetraspanin used commonly as exosome and/or small EV marker, behaved like CD9 rather than like CD63. The novel markers and molecular mechanisms specific of exosomes versus ectosomes identified here will pave the way for further studies to decipher their respective functions.

## RESULTS

### CD63 and CD9 are present on two distinct and one common populations of EVs

In dendritic cells, CD63 and CD9 are secreted abundantly in small EVs, where an EV subpopulation bearing CD63 with CD9 and/or CD81 seemed to correspond to endosome-derived exosomes, but they are also detected in larger ones^7^. We first analyzed their distribution in EVs released by HeLa cells. Both CD9 and CD63 can be found in HeLa EVs isolated by differential centrifugation in the small EV pellet centrifuged at 100Kxg (100K). By centrifuging further at 200Kxg (200K), we recovered some more CD9 and CD63 in material that had not been pelleted at 100K (figure 1A), whereas hardly any signal was detected in the large EVs pelleted at 2Kxg (2K) and 10Kxg (10K). Therefore, CD63 and CD9 are secreted by HeLa mainly in small EVs. For subsequent isolations of EVs by differential ultracentrifugation, we decided to increase recovery by centrifuging the conditioned medium at 200K instead of 100K. To determine whether CD9 and CD63 are on the same or on different sEVs, we immunoisolated these EVs from HeLa concentrated conditioned medium with antibodies specific for one or the other tetraspanin (figure 1B). Side-by-side analysis of the isolated EVs (pull-down: PD) and the EVs that had not bound the antibody (Flow-Through: FT) shows that anti-CD63 co-precipitates around 50% of CD9, and that conversely anti-CD9 co-precipitates around 50% of CD63 (figure 1B). Immunoprecipitation of CD63 and CD9 in a mixed conditioned medium from CD63-KO and CD9-KO cell lines shows no co-isolation of the other tetraspanin by either antibody (supp figure 1). Therefore, the observed co-precipitation of CD63 and CD9 from WT conditioned medium is not due to aggregation of single positive EVs but rather to simultaneous presence of the two tetraspanins on the same EV. This shows that HeLa cells secrete at least three EV populations defined by these markers: one population with both CD63 and CD9, one with CD63 only and one with CD9 only. We next questioned the sub-cellular origin of the CD63+/CD9+ EV population released by HeLa cells. By immunofluorescence, we observed that CD63 is located mainly in intracellular compartments. In contrast, CD9 is mainly found at the plasma membrane but also in rare dim intracellular compartments. No clear co-localization of the two proteins was observed a steady-state (figure 1C). Thus, following the dynamics of CD63 and CD9 localization in the cell during their transport is required to understand biogenesis of the double-positive EVs.

**Figure 1:**
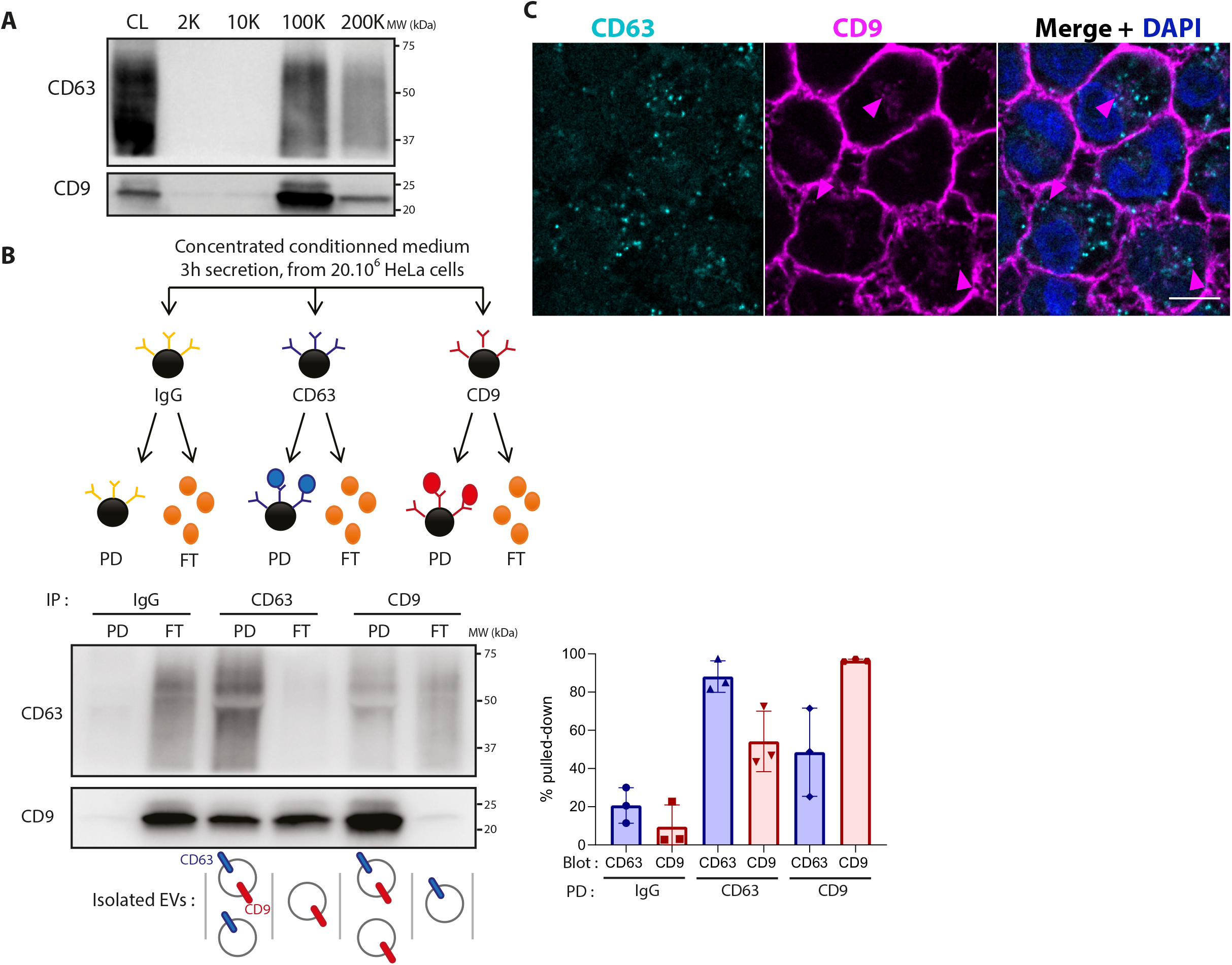
CD63 and CD9 are found on two different and one common EV populations. **A)** Western blot showing CD9 and CD63 detection in cell lysates (CL) and the pellets obtained from HeLa conditioned media after differential ultracentrifugation (2K, 10K, 100K, 200K). The loaded material comes from 20×10^6^ cells for the centrifugation pellets, and from 0,2×10^6^ cells for the cell lysate. Representative of two independent experiments. **B)** Principle of immunoprecipitation of CD63 and CD9 EVs in HeLa concentrated conditioned medium and representative Western blot of the pull-down (PD) and flow-through (FT) of the immunoprecipitation, with quantification of the relative CD63 and CD9 bands intensity of three independent experiments. **C)** Confocal imaging of immunofluorescence staining of CD63 and CD9 in HeLa cells. Pink arrows show CD9 localized in intracellular compartments. Scale bar: 5μm

### CD63 and CD9 traffic transiently through late endosomes and the PM

The RUSH system^8^ was used to follow CD9 and CD63 synchronously by live imaging from the ER to their residency compartment (figure 2A). The principle of this assay is to reversibly retain a protein of interest in a donor compartment, like the ER, and to monitor its release by live imaging. To this end, the protein of interest is fused to a fluorescent moiety and to the streptavidin-binding peptide (SBP), and co-expressed with another protein localized in the donor compartment, fused to streptavidin and used as a hook (here: Streptavidin-KDEL for retention in the ER). Upon synthesis, the SBP-fluorescent protein is retained in the ER by its interaction with streptavidin-KDEL. Biotin is then added to release the protein of interest. Biotin indeed binds to streptavidin and induces the release of the SBP-fused protein that can then follow its normal trafficking route. The fluorescent protein and the SBP were inserted in the small extracellular loop of CD63 and CD9 (figure 2A), following a strategy previously used to follow trafficking of MVBs and their fusion with the PM^9^. Indeed, the CD63-pHLuorin construct used in this previous study behaved similarly to the endogenous CD63 in terms of steady-state intracellular localization and release in EVs, thus showing that this particular site of insertion of the fluorescent protein did not create artefacts for further study of its trafficking. The trafficking of the two tetraspanins was quantified and compared in HeLa transiently cotransfected with the RUSH plasmids for CD63-mCherry and CD9-eGFP (figure 2B). Between 15 and 40 minutes after leaving the ER, both enter and leave the Golgi with similar kinetics. Then they separate (evidenced by a decrease of their Pearson’s co-localization coefficient) and CD63 eventually accumulates in intracellular compartments while CD9 reaches the cell periphery. However, between 30 min and 1h after biotin addition, after leaving the Golgi, a fraction of both proteins was found in some common internal compartments (figure 2B, white arrowheads). Adding NH_4_Cl, which neutralizes the pH of acidic compartments, to cells transfected with the RUSH plasmids for either CD63-eGFP or CD9-eGFP led to an increase of fluorescence, stronger for CD63 than for CD9 (supp figure 2A). This is likely due to unquenching of eGFP which is sensitive to acidic conditions, and this demonstrates that CD63 is going mainly into acidic compartments, but that a fraction of CD9 might also be addressed in such compartments. Immunostaining and electron-microscopy (EM) of eGFP and mCherry in RUSH-CD9 and -CD63 co-transfected cells (figure 2C and supp table 1) confirms that, at 1h after biotin addition, multivesicular bodies (MVBs) containing both CD63 and CD9 can be observed (figure 2C), and CD9 is observed with a Relative Labeling Index (RLI) in MVBs >1 (5.7 in Supp table 1) showing a specific location in these compartments. At 2h, the RLI of CD9 in MVBs decreases to 3.1 but is still >1 (Supp table 1), and these compartments also contain CD63 (figure 2C). At all time points, we observed that a majority of MVBs and lysosomes contained only CD63 and not CD9. The partial co-localization of CD63 and CD9 in some late Rab7-positive endosomes, was confirmed by co-transfecting HeLa with CD9-mCherry and CD63-mTurquoise2 RUSH plasmids and a Rab7-eGFP plasmid (figure 2D). In addition, immuno-EM also showed that CD63 partially and transiently localized at the plasma membrane 1h after biotin addition (figure 2C and supp table 1: RLI = 3.5 at 1h, 1.5 at 2h). Some colocalization of the CD63-RUSH construct with MyrPalm-mCherry, which labels the plasma membrane, was also observed between 20 and 60 min post-biotin addition (supp figure 2B). Two transient places of co-localization of CD63 and CD9 were thus detected: in late endosomes and at the PM.

**Figure 2:**
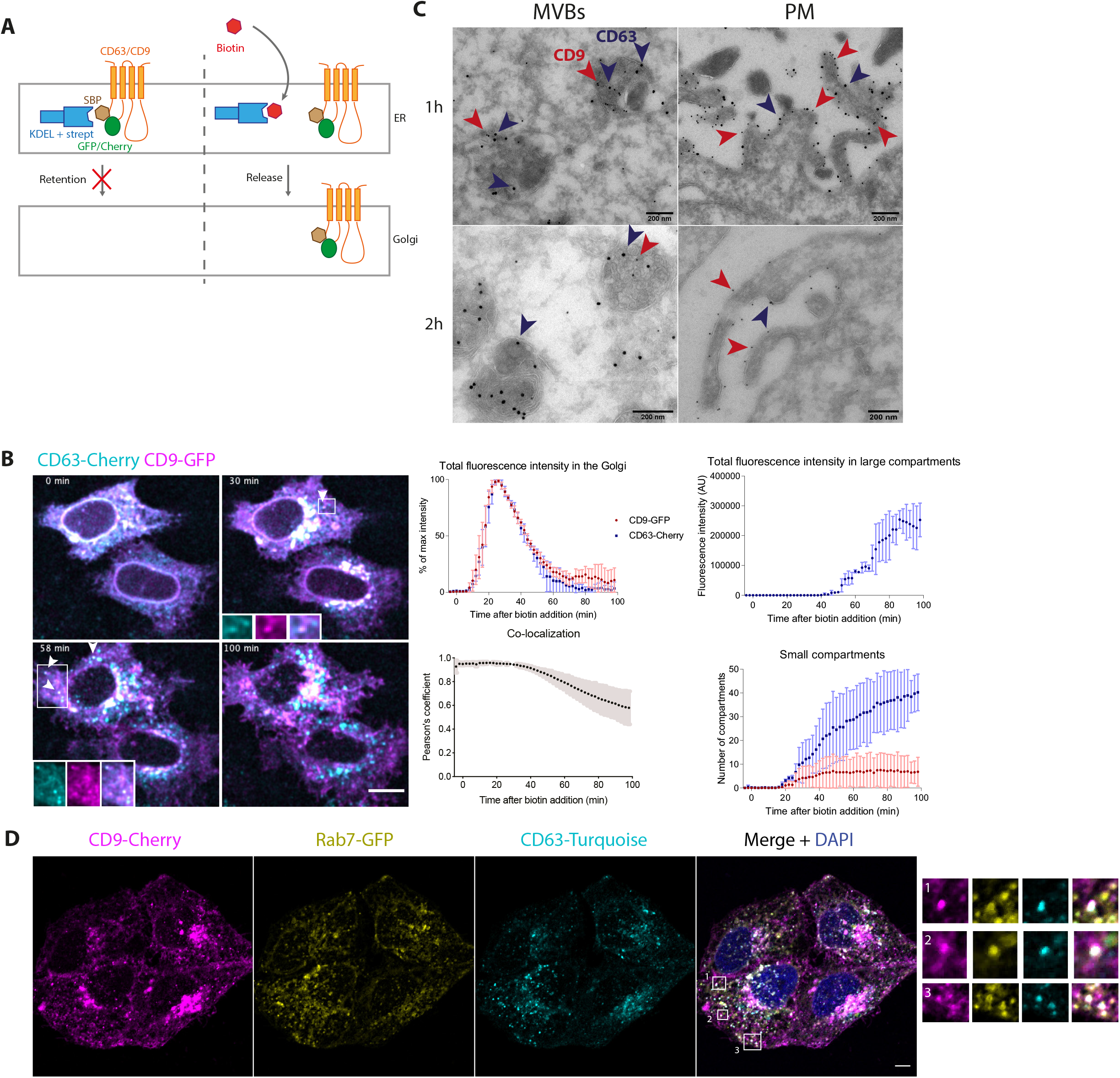
CD63 and CD9 transiently co-localize in multivesicular bodies and at the plasma membrane. **A)** Principle of the RUSH system used to follow CD63 and CD9 intracellular trafficking. SBP: streptavidin binding peptide; strept: streptavidin; ER: endoplasmic reticulum. **B)** Micrographs and quantifications of live imaging of HeLa cells co-transfected with the CD63-mCherry and CD9-eGFP RUSH plasmids. Biotin at 40 μM was added at T=0. White arrows show peripheral compartments where CD63 and CD9 co-localize. Zprojection of 11 planes. Scale bar: 5μm. Quantification upon time of three independent experiments showing the eGFP and mCherry fluorescence intensity in the Golgi and in large compartments, the number of eGFP- or mCherry-positive small compartments and the Pearson’s co-localization coefficient between eGFP and mCherry where automatically quantified. **C)** Representative electron microscopy images of HeLa cells co-transfected with RUSH constructs of CD63-mCherry and CD9-eGFP after 1h or 2h of incubation with biotin, labelled with anti-eGFP gold 10nm (red arrows) and anti-mCherry gold 15nm (blue arrows). Quantification in 7 different fields of the distribution of CD9-eGFP and CD63-mCherry in ER-Golgi, MVBs, on the PM or in residual compartments is provided in Supp table 1. **D)** Confocal microscopy picture of HeLa cells co-transfected with CD63-mTurquoise2 and CD9-mCherry RUSH plasmids and Rab7-eGFP plasmid after 1h of incubation with biotin. Scale bar: 5μm.

### The mutant CD63-YA does not traffic to acidic internal compartments

Since CD63 can be present at the PM, we then asked if it could be secreted in EVs from this location. To answer this question, we first generated a form mutated in its lysosome targeting motif: CD63-YA (GYEVM -> GAEVM) (figure 3A). This mutant has previously been described to interact neither with AP2 nor with AP3, which leads to its plasma membrane localization^10^. Imaging transfected HeLa with the CD63-WT-eGFP or CD63-YA-eGFP RUSH plasmids confirmed that at steady state, contrary to CD63-WT, CD63-YA was located mainly at the cell periphery and small intracellular compartments (figure 3B). CD63-YA arrival and exit from the Golgi showed similar kinetics as those of CD63-WT and CD9 (supp figure 3A). CD63-YA then appeared in small intracellular compartments, rather than large ones like CD63 (supp Figure 3A). When NH_4_Cl was added 1h after biotin addition, only a slight increase of total fluorescence intensity was observed (supp figure 3B), similar to the increase observed for CD9-eGFP (supp Figure 2A). This confirms that CD63-YA does not go to acidic compartments as much as CD63-WT. Finally, co-transfecting HeLa with CD63-YA and CD63-WT or CD9 RUSH plasmids showed a similar trafficking for CD63-YA and CD9 with mainly peripheral localization 2h after biotin addition (figure 3C).

**Figure 3:**
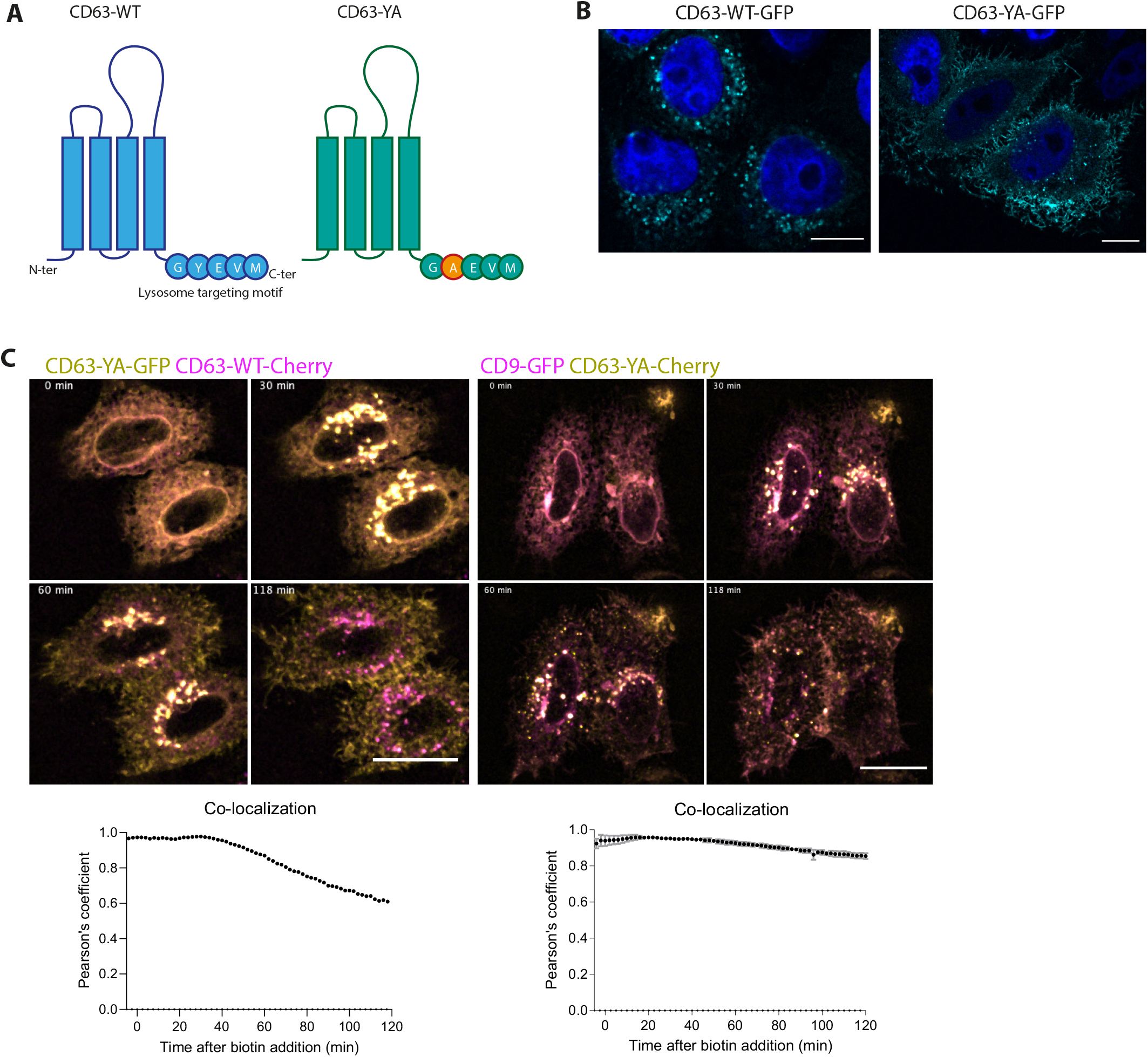
The mutant CD63-YA displays different trafficking from CD63-WT and similar trafficking as CD9. **A)** Scheme of the structure and C-terminal sequences of CD63-WT and the mutant CD63-YA. **B)** Immunofluorescence of HeLa cells transfected with the RUSH CD63-eGFP or CD63-YA-eGFP plasmids at steady state. Scale bar 10μm. **C)** Micrographs of HeLa cells co-transfected with CD63-YA-eGFP and CD63-WT-mCherry or CD9-eGFP and CD63-YA-mCherry, and quantification of the Pearson’s co-localization coefficient over time between eGFP and mCherry. Biotin was added at T=0. Zprojection of 11 planes. Scale bar 5μm. CD63/CD63-YA: n=1 experiment, CD9/CD63-YA: n=2 independent experiments.

### CD9 and CD63-YA have similar kinetics of PM localization and internalization, which differ from those of CD63

To confirm that the different tetraspanins use the PM as transport intermediate, and quantify more precisely the extent and kinetics of such transport, we quantified PM exposure carrying out anti-GFP staining of non-permeabilized HeLa transfected with CD63-WT-eGFP, CD63-YA-eGFP or CD9-eGFP RUSH plasmids. The relative expression of GFP at the surface (anti-GFP-AF647 signal normalized to total GFP expression in fixed cells) was then quantified by flow cytometry at different times after biotin addition and at steady-state corresponding to continuous presence of biotin from the time of transfection (figure 4A). CD9 and CD63-YA reached a similar surface (AF647)/total (GFP) fluorescence intensity ratio of 1 around 1h after biotin addition, which increased further to around 2 at steady-state. For CD63, by contrast, the maximum surface/total ratio increased but remained below 1 until 2h post-biotin addition, and decreased between 2h and the steady-state (figure 4A). This kinetic quantitative analysis thus confirms the microscopy observations: a portion of CD63 is transiently localized at the plasma membrane, whereas CD63-YA and CD9 behave similarly and accumulate at the plasma membrane. These different behaviors could be due to different rates of internalization from the plasma membrane. To quantify this internalization, we performed an antibody uptake assay: 2h after biotin addition, cells were incubated with anti-GFP-AF647 antibodies to label RUSH-tetraspanins at the surface, and incubated at 37°C for 1h. Cells were then treated by trypsin to remove antibodies still present at the cell surface (= stripping, figure 4B) before quantitative analysis by flow cytometry. This treatment efficiently removed antibodies remaining at the cell surface, since the signal was reduced by 77%+/-2,8 for CD63-GFP, 87%+/-0,4 for CD63-YA-GFP and 94,9%+/-0,8 for CD9 in cells incubated at 4°C. While around 80% of CD63 is internalized (i.e. the AF647 signal is reduced by 20% in the stripped condition), only 30% of CD9 and 40% of CD63-YA are internalized (figure 4B). Collectively, the live tracking and cell surface arrival experiments show that CD63-YA behaves mostly like CD9: it accumulates at the cell surface, but a minor portion also undergoes re-internalization, whereas the WT CD63 is quickly and massively re-internalized, as previously shown^10^.

**Figure 4:**
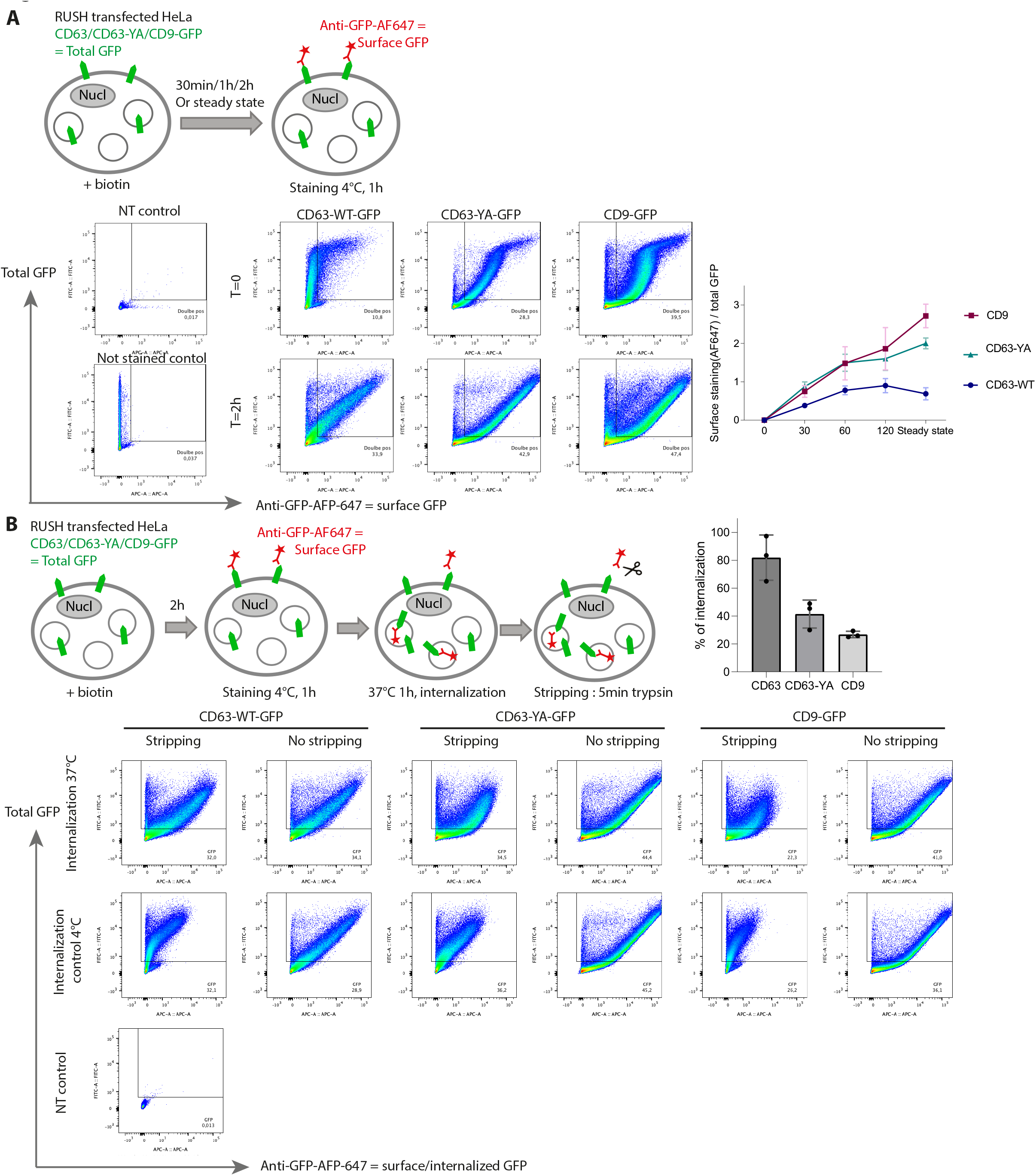
Dynamics of localization at the plasma membrane and of internalization of CD63 are different from those of CD63-YA and CD9. **A)** Principle and flow cytometry analysis of anti-GFP surface staining of HeLa cells transfected with the RUSH constructs CD63-WT-eGFP, CD63-YA-eGFP or CD9-eGFP after different incubation times with biotin followed by fixation. The ratio of the surface staining (AF647) over the total GFP signal mean fluorescence intensities is represented at different time points, time 0 subtracted, for 3 independent experiments. **B)** Principle and flow cytometry analysis of anti-GFP uptake after surface staining of HeLa cells transfected with the RUSH constructs CD63-WT-eGFP, CD63-YA-eGFP or CD9-eGFP after 2h of incubation with biotin. The percentage of internalized anti-GFP-AFP647 is represented for 3 independent experiments.

### EVs released by HeLa contain more CD9 and CD63-YA than CD63

We then asked whether altering subcellular localization of the same molecule, CD63, affects its release in EVs. After 24h of treatment with biotin, the transfected RUSH CD63-YA-mCherry was efficiently released in small EVs (200K pellet, figure 5A). Interestingly, around 50% more mCherry signal was detected for CD63-YA than for CD63-WT. This difference was even more striking when EVs released during 24h with biotin were recovered by anti-GFP immuno-precipitation from HeLa cells transfected with CD9-, CD63-WT- or CD63-YA-eGFP RUSH constructs (figure 5B). While similar numbers of cells expressed the 3 GFP-constructs (legend figure 5B), and similar amounts of EVs were released by the RUSH constructs-transfected cells (supp figure 4A), between 2 and 4 times more GFP signal was detected per particle for both CD63-YA and CD9 than for CD63-WT (figure 5B). BafilomycinA1 (BafA1) is a vacuolar ATPase inhibitor which inhibits the acidification of late endosomes and has been described to stimulate the secretion of CD63+ EVs^11,12^ and we confirmed that CD63-eGFP release in EVs was increased by BafA1. In contrast, no increase was observed for CD63-YA-eGFP or CD9-eGFP upon BafA1 treatment (figure 5C). These results show that CD63 can be efficiently released in small EVs when its location is stabilized at the plasma membrane through mutation of its endocytosis signal, and confirm that different mechanisms are responsible for the release of CD63 in EVs depending on its localization in endosomes or at the PM. They also suggest more abundant release of CD9 and the PM-localized mutant CD63 in EVs than the MVB-enriched WT CD63. Consistent with this hypothesis, in the non-transfected HeLa cells, although both CD9 and CD63 are detected in EV pellets, the ratio of total signal in pellets versus whole cell lysates was always lower for CD63 than for CD9, suggesting also a lower efficiency of release of CD63-bearing EVs (figure 5D). Collectively, these results suggest that in HeLa cells, sEVs are more likely to be secreted from the PM than from MVB under steady-state conditions.

**Figure 5:**
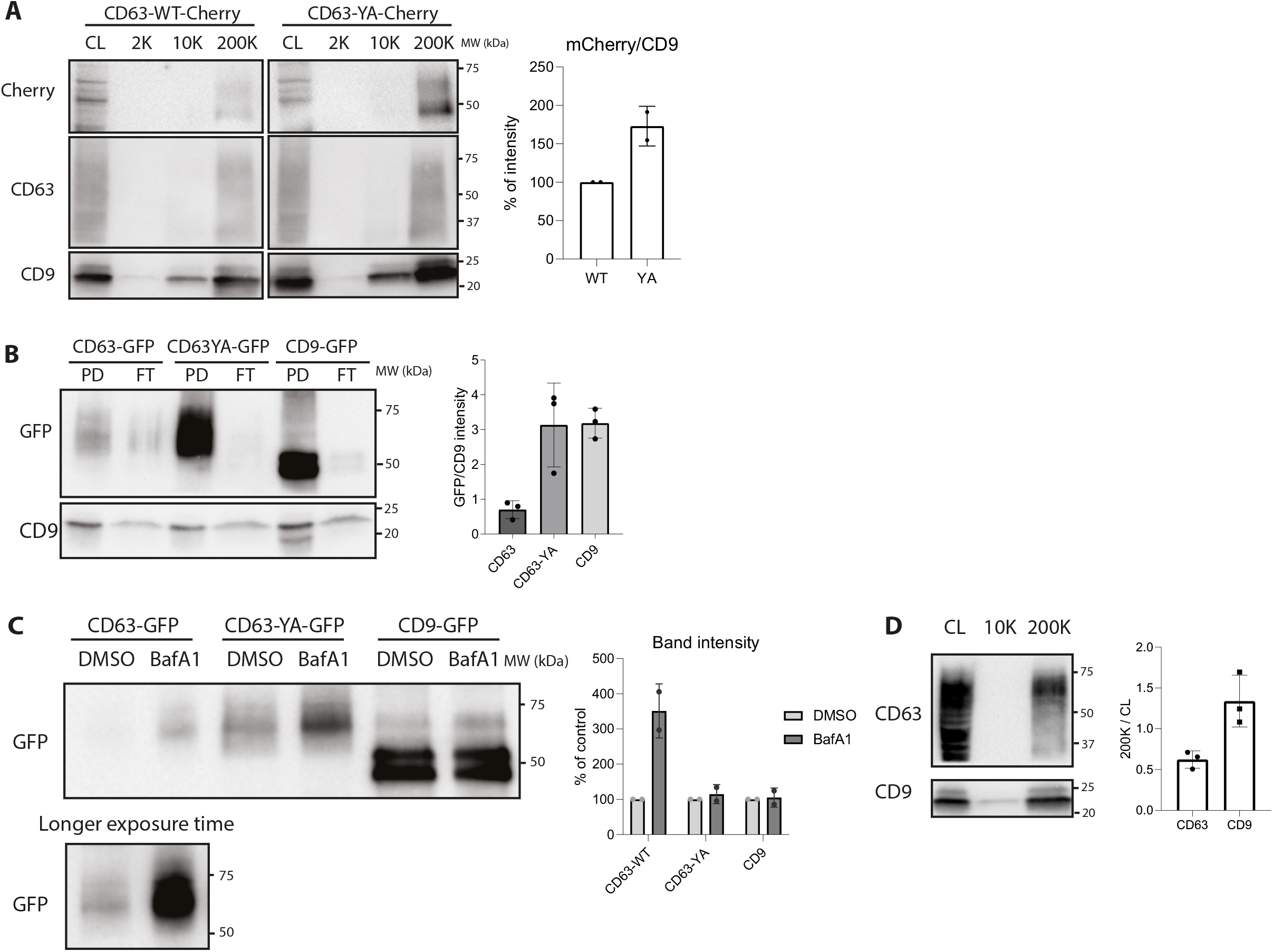
CD63-WT is less efficiently released in EVs than CD9 and CD63-YA. **A)** Western blot of the cell lysate (CL) and of the different EV pellets obtained by differential ultracentrifugation of CCM from HeLa cells transfected with the RUSH plasmids CD63-WT-eGFP or CD63-YA-mCherry (24h release with biotin). EVs from 20.10^6^ cells and CL from 0,2.10^6^ cells were loaded. The intensity of the band corresponding to the mCherry fusion proteins was quantified and normalized by the intensity of the CD9 band and the fold change between CD63-WT and CD63-YA is represented for two independent experiments. **B)** Representative western blot of the pull-down (PD) and flow-through (FT) of the immunoprecipitation of EVs from HeLa cells transfected with the RUSH plasmids CD63-WT-eGFP, CD63-YA-eGFP or CD9-eGFP, recovered 24h after biotin addition. 60.10^8^ total particles quantified by NTA were used for each IP. Percent of GFP+ cells quantified by flow cytometry were similar in the three conditions (respectively 51%±7,6, 47%±6,6, 40%±11). The GFP bands intensity in the PD normalized to endogenous CD9 in the corresponding PD are represented for 3 independent experiments. **C)** Representative Western blot of EVs from HeLa cells transfected with the CD63-WT, CD63-YA or CD9-eGFP RUSH plasmids treated with DMSO of BafA1 100nM during 16h. The same number of EVs between the DMSO and the BafA1 conditions were loaded on the gel (around 100.10^8^ particles). The fold change between DMSO and BafA1 treatment for each construct is represented for two independent experiments. The vacuolar ATPAse inhibitor BafA1 increases specifically EV release of CD63-WT-eGFP. **D)** Proportion of cellular endogenous CD9 and CD63 released in EVs, as semi-quantified on Western blots. The signal for CD9 and CD63 in 200Kg pellets released by 20×10^6^ HeLa cells was divided by the signal for the same molecule in the total lysate of 0.2×10^6^ cells, run on the same blot. 1 representative Western blot and quantification of 3 independent experiments.

### Quantitative mass spectrometry analysis of synchronously-secreted CD63 or CD9 EVs reveals signatures of plasma membrane and endosome derived EVs

To better determine the origin of CD63+ and CD9+ EVs and to identify specific markers of EVs released from PM or from endosomes, we performed a quantitative mass spectrometry analysis of GFP+ EVs recovered by immunoprecipitation from HeLa transfected with the CD63-WT-eGFP or CD9-eGFP RUSH plasmids, incubated during 3h or 24h with biotin. The RUSH system had several unique advantages in this approach. First, the analysis of EVs after short-term trafficking of the RUSH constructs enabled to study a majority of freshly secreted EVs avoiding too many cycles of re-uptake and recycling. Second, use of the same anti-eGFP antibody to isolate both RUSH construct-bearing EVs ensured identical efficacies of immunoisolation, which is necessary for reliable quantitative comparison (and difficult to achieve when using different antibodies, e.g. specific for either CD9 or CD63). Third, specificity of release of the identified proteins with CD63 or CD9 was verified by simultaneously analyzing immunoprecipitated negative control samples containing EVs released by non-transfected cells. A total of 333 and 397 proteins specifically isolated by the anti-GFP IP were quantitatively compared between CD63-eGFP and/or CD9-eGFP EVs at, respectively, 3h and 24h (Supp table 2, Supp table 3 and Supp figure 4B). They were categorized as enriched (over two fold) or unique in CD63- or CD9-bearing EVs, versus common to CD63- and CD9-bearing EVs (see Materials and Methods for detailed criteria). At 3h, 44% of the proteins were common to CD63- and CD9-bearing EVs, 37% were enriched or specific of CD63-EVs and 19% were specific of CD9-bearing EVs. At 24h, slightly more (48%) of the proteins were common to CD63- and CD9-bearing EVs, and slightly less (32%) were specific of CD63-EVs, while the proportion or proteins specific of CD9-EVs remained the same (20%) (figure 6A and Supp figure 4B). This suggests that a common mechanism of release (or subcellular origin) of CD63 EVs and a fraction of CD9 EVs and another mechanism specific of CD63 EVs are both equally important early after CD63 and CD9 synthesis. In addition, there is also a distinct mechanism of secretion of CD9 EVs. Later on, EVs released by the shared mechanism become predominant, which can be due to either more release, or less recapture, or the CD9/CD63 distribution being more ‘equilibrated’ over the various compartments of residence.

**Figure 6:**
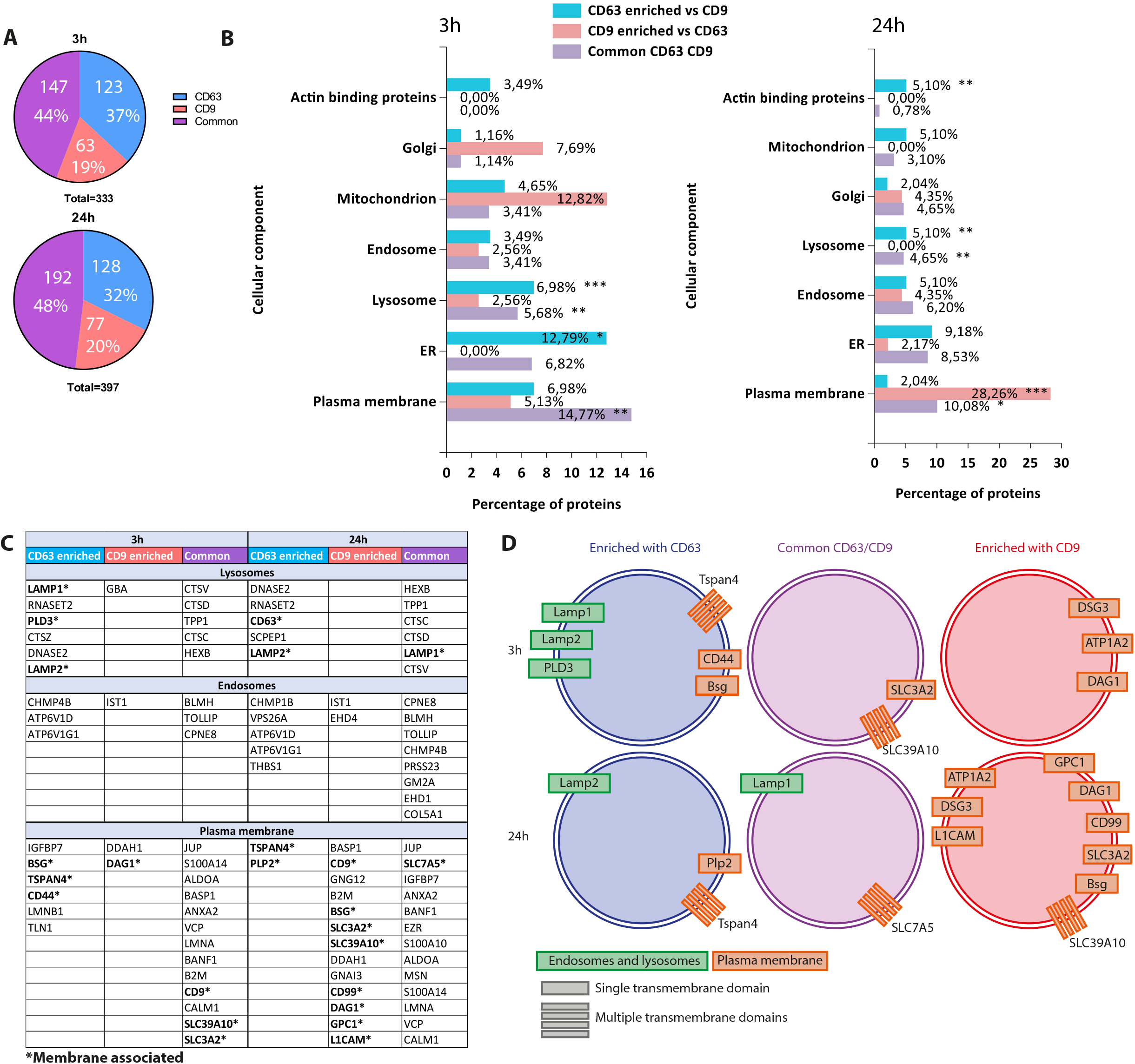
Quantitative proteomic analysis of CD63-eGFP-RUSH and CD9-eGFP-RUSH EVs. EVs were isolated by anti-GFP immuno-isolation from either non-transfected HeLa cells, or HeLa transfected with CD63-eGFP-RUSH or CD9-eGFP-RUSH, either 3h or 24h after biotin addition, and their composition was analyzed by mass-spectrometry. **A)** Pie-charts of number and % of identified proteins in each category (enriched with CD63, or with CD9, or common to both) **B,C**) Results of the FunRich gene enrichment analysis among the proteins either enriched in the CD63-(blue), or CD9-eGFP (red) samples, or common between the CD63 and CD9-eGFP samples (purple) at 3h or 24h after biotin addition. B) For each subcellular compartment gene list, % enrichment represents the % of genes of this category in the list, and the p-value shows the significance of this percentage according to the representation of this category in the database. P-value (hypergeometric uncorrected): *: p < 0,05 ; **: p < 0,01 ; ***: p < 0,001. C) Table showing the names of proteins identified in EVs and used for the endosome, lysosome and PM (plasma membrane) categories are listed. **D)** Schematic representation of the transmembrane proteins identified in the CD63-, CD9-, or CD63/CD9-eGFP EVs at 3h and 24h.

To try to determine the subcellular origin of these EVs, we compared our protein lists with a reference database of >5000 proteins specifically assigned to HeLa subcellular compartments^13^ using the FunRich software^14,15^. For each category of proteins (enriched in CD63-EVs, enriched in CD9-EVs or common to CD63- and CD9-EVs), we calculated the percent assigned to the different intracellular organelles and whether this percent was significantly different with that of the reference database. At 3h (figure 6B, left), the CD63/CD9 common category was significantly enriched for PM proteins (14.8% versus 9.7% in the total proteome), whereas this enrichment was not observed for either CD63- or CD9-EV-specific proteins (7 and 5%, respectively). A significant enrichment was observed for lysosomal proteins in both CD63-EVs and in the CD63/CD9 common category (7 and 5.7% versus 1.7% in the total proteome). Interestingly, transmembrane lysosomal proteins (LAMP1, LAMP2, PLD3) were only found in CD63-EVs, whereas only soluble lysosomal enzymes (CTSV/D/C and TPP1) were enriched in the CD63/CD9 common category (figure 6C). This suggests that a specific sorting of transmembrane lysosomal cargoes into EVs by CD63 may occur, without involvement of CD9, whereas the CD63- and/or CD9-EVs trafficking into lysosomes would capture on their surface some luminal proteins. At 24h (figure 6B, right), we observed in CD9-EVs a strong and significant enrichment of PM components (28.3%), whereas CD63-EVs contained low amounts of PM components (2%). Lysosomal protein enrichment decreased slightly but was still significant in both the CD63-EVs and the CD63/CD9-common category, and LAMP1 was enriched in the latter (figure 6C-D). Thus in 24h conditioned media, a time routinely used to recover EVs, CD9-bearing EVs come from the PM, whereas CD63/CD9-EVs come from both PM and lysosomes.

We then searched for transmembrane (TM) proteins, using both the GO term analysis and manual annotation, which could associate specifically to one or the other EV type and could be used to label or isolate them by antibodies recognizing their extracellular domains. Only 6-14 TM proteins were present in the CD9-EVs (respectively 3h-24h), 8-8 in the common CD63/CD9-category, and 18-16 in the CD63-EVs (red text in Supp table 3). Among these, proteins assigned to lysosomes, endosomes or the PM are shown in figures 6C-D and supp figure 4B. Only PM proteins were enriched in CD9-EVs, whereas both endosome/lysosome and PM transmembrane proteins were enriched in CD63-EVs.

### BafA1 and GW4869 drugs affect differently secretion of CD63 and LAMP1, versus CD9, BSG and SLC3A2

We selected for further analysis: LAMP1 (or CD107a) for its known lysosomal steady-state localization and its enrichment in CD63-EVs and/or in the common CD63/CD9 category, basigin (BSG, a plasma membrane protein also called EMPRINN or CD147) for its different association to CD63 at 3h and CD9 at 24h, and SLC3A2 (a PM channel also called 4F2 cell surface antigen or CD98), for its common presence in CD63- and CD9-EVs at 3h and its enrichment in CD9-EVs at 24h. In addition, we observed an enrichment of syntenin-1 (or SDCBP) in CD63-EVs at 3h, whereas at 24h it was in the common CD63/CD9 category. This protein, described to play a role in exosomes formation^16^, and often used as a small EV marker, was also chosen for further analysis.

We studied the effects of two drugs used in the literature to disturb exosome secretion (BafA1^11,12^ and the inhibitor of neutral sphingomyelinase GW4869^17^) on the secretion of our selected proteins and other sEV markers (syntenin-1/*SDCBP* and CD81). GW4869 treatment decreased more than twice the total number of released particles quantified by Nanoparticle tracking analysis (figure 7A, Log2(fold change)<-1). This was associated with a milder decrease of CD63, CD81 and LAMP1 in EVs (figure 7B right panel, Log2(fold change)>-1), but had no effect on the release of CD9 and SLC3A2 (figure 7B). In contrast, GW4869 decreased more than 4 fold the amount of syntenin-1, and with variable amplitude the amount of BSG (between 0 and 32 times less) in the remaining particles. Therefore, GW4869 inhibits release in EVs of syntenin-1, variably BSG, but not the other proteins. Interestingly, the effect of GW4869 on the total particles release is higher than its effect on most of the analyzed EV markers, suggesting that it affects also a population of EVs of unknown origin (or possibly of non-vesicular particles) carrying syntenin and other markers. On the other hand, the effect of BafA1 was more clear-cut. HeLa cells exposed for 16h to BafA1 released twice as many particles (figure 7A). Molecularly, it resulted in the release of 4 times or more CD63, syntenin-1 and LAMP1, but less than 2 folds more CD9 and CD81, and no effect or a decrease in SLC3A2 or BSG as compared to control cells (measured by Western blot, figure 7B). Together with the specific effect of BafA1 on the release of CD63 and not of CD63-YA (figure 5C), these results thus confirm that the CD63- and LAMP1-positive subpopulation of EVs represent MVB-derived exosomes, which also contain syntenin-1. In contrast, BSG- and SLC3A2-containing EVs are released primarily from the PM. Of note, CD81 like CD9 was only slightly increased by BafA1-mediated endosome alkalization, suggesting that, like CD9, CD81 buds mainly in small ectosomes from the PM. In fact, we observed, by the RUSH approach, a similar trafficking pattern for CD9 and CD81, with final colocalization at the PM (supp figure 5). The BafA1 treatment of EV-secreting cells thus represents a convincing way to evaluate the association of given EV markers to exosomes (increased secretion) versus ectosomes (no or minor increase of secretion).

**Figure 7 :**
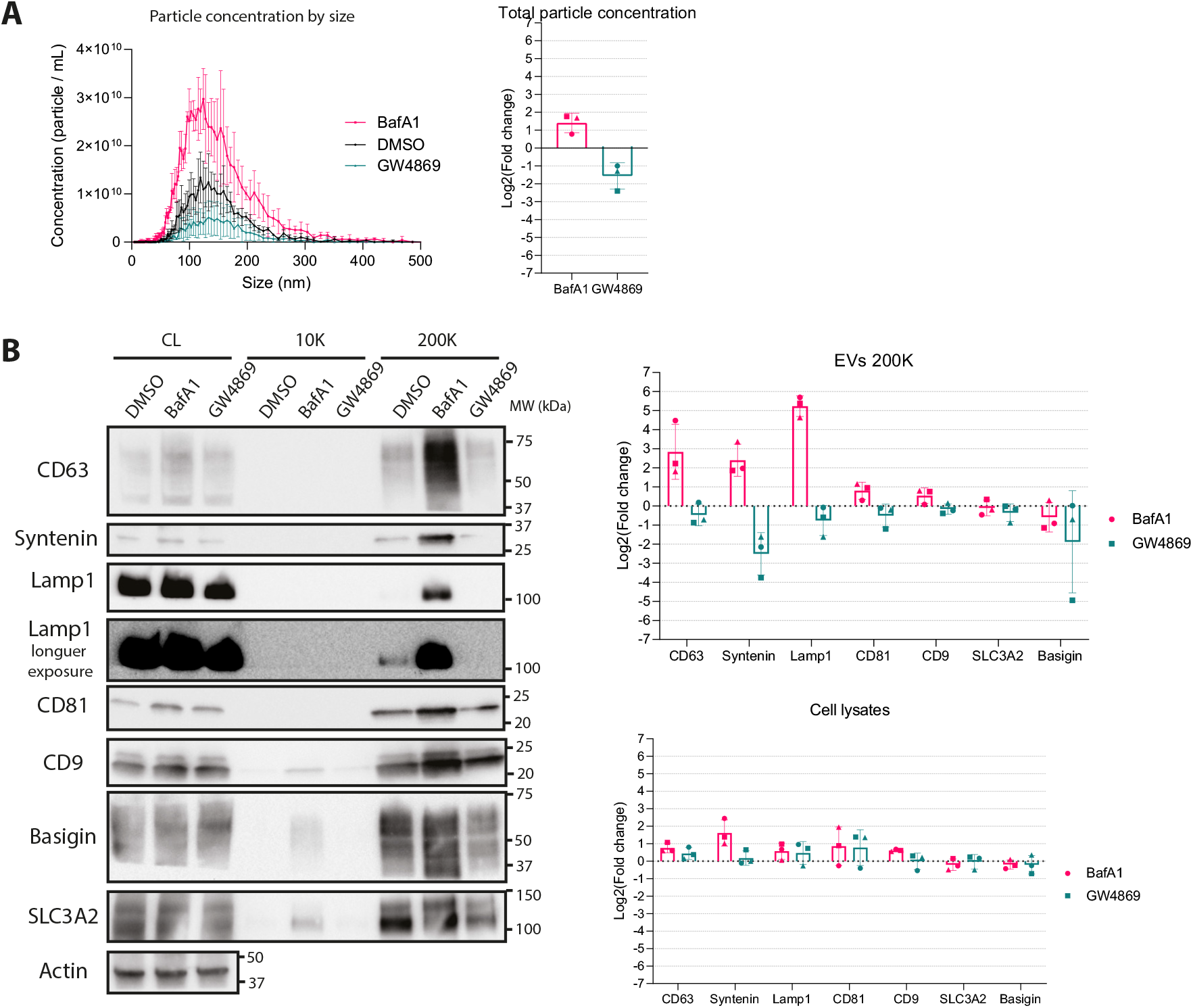
Different effect of BafA1 and GW4869 on secretion of CD63, CD9 and the novel EV markers. **A)** Nanoparticle tracking analysis (NTA) of EVs obtained by differential ultracentrifugation from equal numbers of HeLa cells treated with an equal volume of DMSO (control), Bafilomycin A1 (BafA1) 100nM or GW4869 10μM during 16h. The particles concentration according to their size and the fold change of the total particle concentration between treated and control conditions are represented. **B**) Representative Western blot of cell lysates from 0.2×10^6^ HeLa cells and EVs from 20×10^6^ HeLa cells treated with DMSO, BafA1 or GW4869, corresponding to the samples of A). The fold change between DMSO and BafA1 or GW4869 treatment of the bands intensity in the 200K pellet divided by the cell lysate is represented for three independent experiments.

## DISCUSSION

In this work, we provide evidence that sEVs bearing tetraspanins, especially CD9 and CD81 with little CD63, bud mainly from the plasma membrane, whereas others bearing CD63 with little CD9 but containing some late endosome proteins form in internal compartments and qualify as exosomes. We also identified a small set of additional transmembrane proteins that can be used to distinguish the small ectosomes from *bona fide* exosomes. To obtain these evidences, we followed in live cells the intracellular trafficking of the tetraspanins and identified their colocalization and segregation over time (Figures 2-4), we quantified the release in EVs of an endocytosis-defective mutant form of CD63 which traffics like CD9 (Figures 3-5), we performed quantitative proteomic analysis to identify proteins that are specifically released in EVs with CD9 or CD63 (Figure 6), and we quantified the effect on EV secretion of a drug known to increase the pH of late endosomes, BafA1, to evidence distinct behavior of endosomal versus PM EV markers (Figure 7).

Incidentally, our study revealed that a drug commonly used as an exosome inhibitor, GW4869, which inhibits neutral sphingomyelinase and thus prevents ceramide accumulation, should be used with caution. The low or variable effect of GW4869 on release of our EV markers compared to its effect on the global number of released EVs questions its specificity for exosomes. This drug might affect a specific subpopulation of EVs containing syntenin-1, but none of the other exosome markers in HeLa. In addition, the requirement of ceramide for EV secretion largely depends as well on the cell type. For instance, in the breast carcinoma cell line SKBR3, GW4869 increased release of ectosomes while it decreased the amount of small EVs^18^. We did not observe higher levels of large EVs released by GW4869-treated HeLa cells. Since ceramide accumulation can lead to many additional effects than changing budding at the PM or in endosomes, for instance by promoting apoptosis, interpretation of GW4869 effects of EV release is always difficult.

Comparison of small EVs (containing a mixture of exosomes and small ectosomes), and larger EVs/ectosomes showed differences in their respective protein composition and oncogenic activities^19,20^, but these studies did not explore the diversity within small EVs. More recently, further approaches to separate subtypes of small EVs and compare their cargoes have been published^7,21,22^. Importantly, the two most recent studies demonstrated the presence of non-EV components, called extracellular nanoparticles^22^, and small lipidic structures called exomeres of unclear vesicular nature^21^ within the bulk of small EV preparations. Subtypes of small EVs of different sizes within the 50-150 nm range^21^, or slightly different densities in gradients^7,22^, were also reported. However, the subcellular origin of these different small EVs was not specifically evaluated, and an exosomal nature could only be speculated, based on enriched presence of molecules known to associate with endosomes in some of the recovered small EVs. Our study has the advantage of combining the use of several markers from endosomes and lysosomes or from the PM with a search for mechanisms involved in their secretion by chemical inhibition, to demonstrate the existence of different subpopulations of small EVs, to characterize their protein content and their subcellular origin.

Our results demonstrate that HeLa cells release more small ectosomes than *bona fide* exosomes. An interesting tool that allowed us to demonstrate this proposal is the mutant CD63 molecule devoid of its lysosome-addressing signal, CD63-YA. CD63-YA accumulates at the PM, like CD9, and is more efficiently released in EVs by HeLa cells than the WT CD63 (Figure 5). The eGFP-RUSH constructs of CD63 and CD63-YA could now be used as tools to quantify relative secretion of exosomes versus ectosomes in any cell, by transient transfection followed by quantification of the GFP signal recovered from each construct in the conditioned medium. It would be interesting to analyze this way cells secreting at steady-state more CD63 in EVs than HeLa, and determine whether they indeed display a high ability to secrete *bona fide* exosomes, or whether instead their CD63-EVs mainly form at the PM.

Our results also led us to propose to use a restricted combination of molecules (CD63, LAMP1, syntenin-1, CD9, CD81, BSG and SLC3A2) to determine the proportion of ectosomes versus exosomes in a given small EV preparation. Concerning LAMP1, which behaves like CD63 in response to endosomal pH neutralization, its presence on EVs suggests that exosomes can be released from secretory lysosomes as well as from late endosomes. Concerning the PM-EV markers: SLC3A2 is the heavy chain of various heterodimeric amino-acid transporters, and one of its possible partners, SLC7A5 (also named E16 or LAT-1)^23^, was less specific of the CD9-EVs, but not enriched in the CD63-EVs, thus confirming an ectosomal release. BSG and SLC3A2 directly interact together^24^, and these two proteins are part of a protein complex interacting with integrins^25^. BSG also interacts with CD44^26^: interestingly, BSG and CD44 were specifically present in CD63-bearing EVs at an early time point of secretion, whereas BSG and SLC3A2 were specific to CD9-EVs at a later time point (Figure 6D). thus BSG is probably released with different types of EVs, depending on its interacting partner. Of note, SLC3A2 is highly expressed in several cancers^27^, and BSG has already been described as a cancer cells ectosome marker^18,28^, which makes these two proteins good candidates as tumor cells EVs biomarkers.

In addition, a few other markers identified by the proteomic analysis could be in the future also used as either exosome or ectosome-specific markers (Figure 6). This includes for instance LAMP2, PLD3, PLP2 or the tetraspanin TSPAN4 as exosome markers, and L1CAM, DAG1, CD99 or DSG3 as ectosome markers. Another interesting ectosome marker could be PTGFRN, a known major partner of CD9 and CD81, which has been shown to remain associated with these tetraspanins on EVs secreted by K562 cells^29^: it was identified by 18 peptides with CD9-EVs at 24h (Supp table 2), but could not be reliably compared to CD63-EVs because a single common peptide was present in the two samples. BafA1 treatment of cells in combination with quantitative assessment of the resulting EV level of these EV-associated proteins would have to be performed to validate their proposed specificity. Of course, although the action of BafA1 on EVs seems exosome-specific compared to ectosomes in HeLa according to our observations, this should be further demonstrated in other cell types, by similarly assessing an array of EV-associated proteins. In particular, BafA1 can affect also Golgi trafficking^30^: if prominent over the late endosome-specific effect in some cells, this action of BafA1 would be expected to affect more generally secretion and trafficking of most transmembrane proteins, hence the global composition of EVs. Finally, it would be interesting to use other drugs, recently tested in a small screen specifically designed to quantify CD63-EV release^12^, to quantify their effect on EV release of the new ectosome and exosome markers, and of the mutant CD63-YA reporter.

Our findings and our methodology to discriminate different sEVs subpopulations will be a basis for further studies on the heterogeneity of EVs and their specific functional properties.

## Material and methods

### Cell culture and transfections

HeLa cells were cultured in Dulbecco’s modified Eagle’s medium (DMEM-Glutamax, Gibco), with 10% fœtal calf serum (FCS, Gibco), 100 U/mL penicillin and 100μg/mL streptomycin (Gibco).

HeLa KO for CD63 or CD9 were generated by CRISPR/Cas9 using sgRNA targeting the following sequences :

CD63 CCAGTGGTCATCATCGCAGT and CD9 GAATCGGAGCCATAGTCCAA, which were selected using the CRISPR design tool available at the Broad Institute (https://portals.broadinstitute.org/gpp/public/analysis-tools/sgrna-design). The corresponding guide DNA sequences were cloned into the lentiCRISPRv2 plasmid (Addgene #52961) according to the instructions of the Zhang laboratory (https://www.addgene.org/52961)^31^. The plasmids were transfected using Fugene HD according to the manufacturer’s instructions, and cells were treated after 36-48h with 5μg/ml puromycin for 36-48h. Cells were stained for CD63 or CD9 and negative cells for the antigen of interest were sorted using a FACS Aria cell sorter (Becton Dickinson). CD63 KO cells were subcloned after two sorting rounds and one clone was kept for experiments. CD9 KO cells were subcloned after two sorting rounds and a pool of 5 clones was used for experiments.

Transient plasmids transfections were performed on 80% confluent cells using the calcium phosphate method: 2,5ug/mL of plasmids were mixed 1/20 vol/vol with 1mM Tris pH 8 and 2.5M CaCl_2_ and incubated 5min at room temperature (RT) before being mixed with an equal volume of HEBS (160mM NaCl, 1.5 mM Na_2_ HPO_4_, 50 mM Hepes, pH 7.05). This mix was added at 1/10 vol/vol into culture medium and cells were incubated during 8h before washing with PBS and replacing with fresh culture media for an additional 16h before performing experiments.

### Antibodies and reagents

Primary antibodies for western blot: mouse anti-human CD63 (BD Bioscience clone H5C6), - human CD9 (Millipore clone MM2/57), -GFP (Invitrogen clone GF28R), -human CD147/BSG (Proteintech, clone 1G12B5), -human CD98/SLC3A2 (Proteintech, clone 2B10F5), -human actin (Millipore, clone C4); rabbit anti-human LAMP1 (GeneTex, clone EPR4204), -mCherry (Biovision, polyclonal). Monoclonal rabbit anti-human syntenin was a gift from P. Zimmermann.

Secondary antibodies: HRP-conjugated goat anti-rabbit IgG (H+L) and HRP conjugated goat anti-mouse IgG (H+L) were purchased from Jackson Immuno-Research.

Antibodies for immunoprecipitation: mouse anti-human CD63 (BD Bioscience clone H5C6), mouse anti-human CD9 (Millipore clone MM2/57), normal mouse IgG (Millipore); rabbit anti-GFP polyclonal antibody was purchased from the Tab-IP antibody and the recombinant proteins facilities at the Institut Curie.

Antibodies for immunofluorescence: mouse IgG2b anti-human CD63 (clone TS63b^32^) and mouse IgG1 anti-human CD9 (clone TS9^33^), goat anti-mouse IgG2b Alexa fluor 647 (Invitrogen), goat anti-mouse IgG1 Alexafluor 488 (Invitrogen)

Antibody for surface staining and uptake experiments: mouse anti-GFP-Alexa fluor 647 (BD Biosciences clone 1A12-6-18)

Biotin was purchased from Sigma-Aldrich, a stock solution at 4mM in DMEM was used at 1/100 in culture medium (40μM final) for all RUSH experiments. BafilomycinA1 ready-made solution 0.16mM in DMSO was purchased from Sigma-Aldrich and used at 100nM. GW4869 was purchased from Sigma-Aldrich, dissolved to 5mM in DMSO and used at 10μM. DMSO or drugs were diluted in fresh EV-depleted medium in which cells were cultured for 16h before collection of conditioned medium.

### Plasmids

A RUSH vector encoding for the KDEL-Streptavidin Hook was used for all CD63, CD9 and CD81 reporters. A CD63-SBP-pHluorin RUSH plasmid was initially generated by inserting the SBP sequence upstream of the pHluorin fragment in the CD63-pHluorin sequence described in ^9^. The CD63-SBP-pHluorin fragment was synthesized and ligated in a Str-KDEL Neomycin vector after AscI/PacI digestion, thereby replacing the neomycin cassette with CD63-SBP-pHluorin. The pHluorin sequence was then replaced by the eGFP, the mCherry (fragments taken from plasmids described in ^8^) or the mTurquoise2 sequence (synthesized by Integrated DNA Technologies) to generate RUSH plasmids allowing the expression of human CD63 with the SBP and eGFP, mCherry or mTurquoise2 in the small luminal loop. The RUSH CD9-eGFP plasmid was obtained from this plasmid, replacing the CD63 sequence by a synthetic sequence of human CD9 (N-terminal sequence: ATGcctCCGGTCAAAGGAGGCACCAAGTGCATCAAATACCTGCTGTTCGGATTTAA CTTCATCTTCTGGCTTGCCGGGATTGCTGTCCTTGCCATTGGACTATGGCTCCGAT *TCGAC**gaattc**ACCGGT**ggccggcc**AACCGGTGGAGCTCGAATCAGATCTTCTCAGA…)* containing restriction sites (in bold) allowing the later cloning of the SBP-eGFP or mCherry sequence from the RUSH-CD63 plasmids and a linker (italics) (gblock purchased from Integrated DNA Technologies). The RUSH CD63-YA-mCherry plasmid was obtained by performing directed mutagenesis on the RUSH CD63-mCherry plasmid. The following primers were used for amplification (mutation site highlighted in bold) :

forward: TGAAGAGTATCAGAAGTGGC**GC**CGAGGTGATGTAGttaatT,

reverse: AattaaCTACATCACCTCG**GC**GCCACTTCTGATACTCTTCA.

The parental plasmid was digested with the DpnI enzyme (NEB). The mCherry was then replaced by eGFP to generate the RUSH CD63-YA-eGFP plasmid. The Myr-Palm-mCherry plasmid was generated by fusing the myristoylation-palmitoylation sequence (in bold) and a linker (ATG**GGCTGCATCAAGAGCAAGCGCAAG**GACAACCTGAACGACGACGGCGTGGACgaaccggtcgccacc) in N-term of the mCherry sequence. The Rab7-eGFP was a gift from P. De Camilli.

### EV isolation by differential ultracentrifugation

FCS-EV depleted medium was first prepared by centrifuging DMEM with 20% FCS at 200Kg overnight with a 45Ti rotor (Beckman Coulter). Supernatant was then recovered by pipetting without taking the last 5mL to avoid disturbing the pellet, and filtered through a 0,22um bottle filter (Millipore). Culture medium of (generally) 3 subconfluent (less than 80%) 150cm dishes was changed, after a PBS wash, for FCS-EV depleted media diluted with DMEM to 10% FCS. 24h or 16h later, medium was collected for EV isolation and cells were trypsinized and counted (generally around 25.10^6^ cells/dish). 3 consecutive centrifugations at 300g at 4°C were performed to remove any floating cells. Then serial centrifugations of the supernatant at 2Kg, 10Kg, 100Kg and 200Kg at 4°C were performed. 2Kg centrifugation was performed during 20min, the pellet washed in 50mL PBS and re-centrifuged for 20min at 2Kg. 10Kg, 100Kg and 200Kg centrifugations were performed in a 45Ti rotor (Beckman Coulter) during 40min, 1h30 and 2h respectively. Pellets were resuspended in 6mL PBS for a wash and centrifuged again at the same speed in a MLA80 rotor (Beckman Coulter) during 20min, 50min and 1h10 respectively. Alternatively, for experiments without a 100Kg centrifugation, the 200Kg centrifugation was performed for 2h immediately after the 10Kg one. Finally, the pellets were resuspended in 1uL PBS/ 1.10^6^ secreting cells.

### Western blot

Cell lysates (CL) for western blot were performed by incubating 1.10^6^ cells in 25uL of lysis buffer (50 mM Tris, pH 7.5, 0.15 M NaCl, 1% Triton X-100) with 2% complete protease inhibitor (Roche) for 20min on ice, followed by a 13000rpm centrifugation for 15min at 4°C to recover the supernatant. EV from 20×10^6^ cells (or specified number of particles) and CL from 0.2×10^6^ cells were mixed with Laemmli sample buffer (BioRad), including 10% of β-mercaptoethanol (Sigma) for the samples needing reducing conditions. After boiling 5min at 95°C, samples were loaded on a 4-15% Mini-protean TGX-stain free gels (BioRad). Transfer was performed on Immuno-Blot PVDF membranes (BioRad), with the Trans-blot turbo transfer system (BioRad) during 7min. Blocking was performed during 30min with Roche blotting solution in TBS 0,1% Tween or with 5% milk in PBS 0,1% Tween for the anti-GFP antibody. Primary antibodies were incubated overnight at 4°C and secondary antibodies during 1h at room temperature (RT). Development was performed using either the BM Chemiluminescence Western blotting Substrate (POD) (Roche), Clarity western ECL substrate (BioRad) or the Immobilon Forte Western HRP substrate (Millipore), and the Chemidoc Touch imager (BioRad). Intensity of the bands was quantified using ImageJ.

### Immunoprecipitation for Western blot

For immunoprecipitation, Pierce protein A magnetic beads (ThermoFisher) were incubated with IgG, CD63 or CD9 antibodies overnight at 4°C on agitation. Concentrated conditioned medium (CCM) was prepared using serum-free DMEM incubated with secreting cells during 3h. This medium was centrifuged at 300g for 10min and at 10Kxg for 40min to remove floating cells and the largest EVs and filtered using Sartorius Vivaspin 100KDa molecular weight cutoff concentrators. On the same day, CCM were incubated overnight with the hybridized beads in PBS 0,001% Tween on agitation at 4°C. Samples were separated from the beads by heating at 95°C during 5min after addition of the loading buffer.

### Immunofluorescence

Cells were seeded on 12mm diameter coverslips. Once they reached confluency, they were fixed with 4% paraformaldehyde (PFA) (EMS) during 15min at RT. Primary and secondary antibodies were successively incubated during 1h each at RT. Coverslips were then mounted on slides with Fluoromount G with DAPI (Invitrogen). Images were acquired on a Zeiss LSM780 confocal microscope using a 63x objective with 1.4 size aperture and processed with ImageJ.

### Live imaging of RUSH constructs

Cells were transfected 24h before imaging and washed with PBS before adding fresh complete culture medium 8h later. To analyze steady-state distribution of the RUSH constructs, cells were transfected and cultured throughout in the presence of 40μM biotin. For synchronized analysis of RUSH constructs, cells were transfected and cultured in complete medium until the time of biotin addition. Live imaging was performed with an Eclipse 80i microscope (Nikon) equipped with spinning disk confocal head (Yokogawa), using a 60x objective with 1,4 size aperture and a iXon Ultra897 camera (Andor). 25mm-diameter coverslips with the transfected cells were put in a L-shape tubing Chamlide (Live Cell Instrument), filled with pre-warmed carbonate independent Leibovitz’s medium (Invitrogen) with 1% FCS. Medium was replaced by pre-warmed Leibovitz’ medium (Gibco) with 1% FCS and 40μM biotin at time 0. For experiments with NH_4_Cl, medium was replaced after 1h of incubation with biotin for medium with biotin and 50mM NH_4_Cl. For quantifications, automated image analysis was performed with ImageJ. We generated macros to quantify the kinetics of trafficking. Briefly, a first macro was used to count along time the number of small punctual fluorescent compartments (less than 15 pixels), using the “find maxima” function. This homemade macro was also used to measure the fluorescence intensity in the Golgi (defined by a threshold of 20 pixels) and in large compartments (threshold of 15 pixels) when the fluorescence in the Golgi decreased. Starting time for measurement of fluorescence in large compartments after exit from the Golgi was manually defined. Another macro was used for the co-localization analysis. Pearson’s coefficients between the 2 markers were computed inside the cytoplasm at each time using Coloc2 plugin. A Pearson’s coefficient of 1 indicates full colocalization.

### Electron microscopy

Sample preparation, ultrathin cryosectioning and immunolabelling were performed as already described^34^. In brief, HeLa cells were grown on culture dishes and fixed by the addition of freshly prepared 4% PFA in 0.1 M phosphate buffer (pH 7.4) to an equal volume of culture medium for 10 min, followed by postfixation with fresh 4% PFA overnight at 4 °C. After rinsing with PBS, the blocks were embedded in 12% gelatin, cryoprotected with 2.3 M sucrose, and frozen in liquid nitrogen. Ultrathin cryosections were cut on a Leica ultracut UC7 cryomicrotome and picked up in a freshly prepared 1:1 mixture of 2.3 M sucrose and 1.8% methylcellulose, thawed and collected on formvar-coated grids. After washing with PBS containing 0.02 M glycine, sections were incubated with primary antibodies and protein A-gold conjugates (PAG) (Utrecht University, The Netherlands). The following antibodies were used at the indicated dilutions: anti-GFP abcam ab290 1:800; anti mCherry GenTex GTX 128508 1:250. Sections were examined using a Tecnai Spirit electron microscope (FEI Company) equipped with a digital camera Quemesa (SIS). Quantitative IEM analysis was used as described by Mayhew *et al.*^35^. A total of four compartments, including MVBs, ER, Golgi and PM, were selected to analyze the intracellular distribution of CD63 and CD9. The expected distribution was obtained by superimposing to pictures an array of points that was generated digitally and points (P) were counted in the selected compartments. Pictures were taken randomly with the only criterion of a well-preserved morphology, and 7 different pictures were used for quantification. Gold particles on all sampled fields were also counted and named as observed gold particles (Ngo). For each compartment, the observed numbers of golds (Ngo) was compared with the expected numbers of golds (Nge, derived from the observed frequencies of point P). LD is calculated as the number of gold particles per test point (LD = Ngo/P). For each compartment, RLI = LD comp /LD cell. RLI = 1 indicates random labeling but RLI > 1 indicates when compartments are preferentially labeled^35^. By means of a two-sample Chi-squared (χ2) analysis with two columns (Ngo) and (Nge) and c compartments (arranged in rows), two distributions were compared, the total and partial χ2 values were calculated, and whether to accept or reject the null hypothesis was decided (of no difference between distributions) for c-1 degrees of freedom. For any given compartment, the partial χ2 is calculated as (Ngo-Nge)^2^/Nge. If the observed and expected distributions are different, examining the partial χ2 values will identify those compartments that are mainly responsible for that difference. A convenient arbitrary cut-off is a partial χ2 value accounting for 10% or more of total χ2^35^. Complete quantification shown in Supp table 1.

### Surface staining

After 30min, 1h, 2h of incubation with biotin, or no incubation (T=0), or at steady-state (cells transfected already in the presence of biotin), cells were detached with 0,5mM EDTA, washed and resuspended in cold PBS with 1% FCS and placed on ice to stop the trafficking during 10min. Cells were then incubated with the anti-GFP-AF647 for 40min on ice and washed, before fixation with 2% PFA at RT during 10min. After washes, cells were resuspended in PBS with 1% FCS and analyzed for GFP and AF647 fluorescence on a FACS Verse (BD). Nontransfected and non-AF647-stained cells were used for gating the cells positive for GFP and AF647. The increase of surface exposure at a given time point was calculated for cells in this gate by the formula: (MFI_AF647_/MFI_GFP_)tn - (MFI_AF647_/MFI_GFP_)t0.

### Antibody uptake

After 2h of incubation with biotin, cells were 1) washed with cold PBS with 1% FCS and placed on ice for 10min, 2) incubated with anti-GFP-AF647 during 1h on ice and washed, 3) placed into complete culture medium at 37°C or 4°C (negative control condition with no internalization) during 1h, and 4) incubated for 5min with trypsin (stripping) or 0,5mM EDTA (no stripping) at 37°C. Detached cells were then fixed with 2% PFA for 10min at RT, washed, and analyzed with the FACS Verse. Not transfected cells were used for gating the GFP positive cells. The percentage of GFP that has been internalized during 1h at 37°C is calculated from the ratio between the stripped and non-stripped conditions of AF647/GFP signal (in the GFP-positive gate), after subtracting the background internalization signal calculated similarly at 4°C, by the following formula: (MFI_stripping_x100)/MFI_no stripping_)37°C-(MFI_stripping_x100)/MFI_no stripping_)_4°C_

### Proteomics of immunocaptured RUSH CD63- and CD9-eGFP EVs

2,5.10^6^ cells per 10cm dish were plated on the day before transfection. 24h after transfection, DMEM without FCS with 40nM biotin was incubated during 3h or 24h with cells transfected with the RUSH plasmids CD63- or CD9-eGFP, or non-transfected cells as control. Harvested medium from 12 plates per transfection or 18 plates of non-transfected cells was centrifuged 20min at 300g and concentrated with Centricon plus-70 filters (Millipore) with a 100KDa molecular weight cut-off. The CCM concentrated to 500uL was then submitted to size exclusion chromatography using qEV 70nm columns (Izon). Fractions 7 to 12 were recovered and pooled according to manufacturer’s instructions. The pooled fractions were finally concentrated using 500uL 100kDa molecular weight cut-off concentrators (Sartorius). The percentage of transfected cells was determined by flow cytometry and the concentration of recovered EVs was measured by nanoparticle tracking analysis with the ZetaView device (Particle Metrix). For 3h samples, EVs from 20.10^6^ GFP-positive cells were incubated with the beads. An equivalent number of total particles from the non-transfected control sample was used. For 24h samples, a total of 60.10^8^ particles was incubated with the beads. The protein G beads were previously hybridized with the anti-GFP overnight and cross-linked using BS^3^ (ThermoFisher) for 1h at 4°C and finally washed with PBS 0,001% Tween. The incubation with the beads was performed immediately after isolation, in PBS 0,001% Tween overnight at 4°C in rotation. The beads were then washed three time with PBS 0,001% Tween with 300mM NaCl and three times with PBS. The flow through recovered for Western blot was concentrated using 500uL 100kDa molecular weight cut-off concentrators. Immuno-isolated vesicles on beads were eluted with 100 μL 80/20 MeCN/H2O + 0.1 % TFA. Dry pellets were solubilized and reduced in 20 μL 8M urea, 200 mM ammonium bicarbonate, 5 mM dithiothreitol, pH 8 with vortexing at 37°C for 1 h. After cooling to room temperature, cysteines were alkylated by adding 10 mM iodoacetamide for 30 min in the dark. After diluting to 1 M urea with 200 mM ammonium bicarbonate pH 8.0, samples were trypsine/LysC (1 μg, Promega) digested in a total volume of 200 μL with vortexing at 37°C overnight. Samples were then loaded onto homemade C18 StageTips for desalting. Peptides were eluted using 40/60 MeCN/H2O + 0.1% formic acid, vacuum concentrated to dryness and reconstituted in injection buffer (2% MeCN/0.3% TFA) before nano-LC-MS/MS analysis. Four independent biological replicates of each sample were analysed.

### Mass Spectrometry Analysis

Liquid chromatography (LC) was performed with an RSLCnano system (Ultimate 3000, Thermo Scientific) coupled online to a Q Exactive HF-X mass spectrometer (MS) with a Nanospay Flex ion source (Thermo Scientific) peptides were first trapped on a C18 column (75 μm inner diameter × 2 cm; nanoViper Acclaim PepMapTM 100, Thermo Scientific) with buffer A (2/98 MeCN/H2O in 0.1% formic acid) at a flow rate of 2.5 μL/min over 4 min. Separation was then performed on a 50 cm x 75 μm C18 column (nanoViper Acclaim PepMapTM RSLC, 2 μm, 100Å, Thermo Scientific) regulated to a temperature of 50°C with a linear gradient of 2% to 30% buffer B (100% MeCN in 0.1% formic acid) at a flow rate of 300 nL/min over 91 min. MS full scans were performed in the ultrahigh-field Orbitrap mass analyzer in ranges m/z 375–1500 with a resolution of 120 000 at m/z 200. The top 20 intense ions were subjected to Orbitrap for further fragmentation via high energy collision dissociation (HCD) activation and a resolution of 15 000 with the intensity threshold kept at 1.3 × 105. We selected ions with charge state from 2+ to 6+ for screening. Normalized collision energy (NCE) was set at 27 and the dynamic exclusion of 40s. For identification, the data were searched against the Homo sapiens (UP000005640) UniProt database using Sequest-HT through Proteome Discoverer (version 2.4). Enzyme specificity was set to trypsin and a maximum of two-missed cleavage sites was allowed. Oxidized methionine, Carbamidomethyl cysteines and N-terminal acetylation were set as variable modifications. Maximum allowed mass deviation was set to 10 ppm for monoisotopic precursor ions and 0.02 Da for MS/MS peaks. The resulting files were further processed using myProMS^36^ v3.9. FDR calculation used Percolator^37^ and was set to 1% at the peptide level for the whole study. The label free quantification was performed by peptide Extracted Ion Chromatograms (XICs), computed with MassChroQ version 2.2.1^38^. XICs from proteotypic peptides between compared conditions (TopN matching) with missed cleavages were used. Median and scale normalization was applied on the total signal to correct the XICs for each biological replicate (N=4). To estimate the significance of the change in protein abundance, a linear model (adjusted on peptides and biological replicates) based on a two-tailed T-test was performed and p-values were adjusted with the Benjamini–Hochberg FDR procedure. In order to eliminate non-specifically isolated proteins, only proteins with at least two peptides across 2 biological replicates of an experimental condition and that showed a log2(fold change) > 0 in the samples containing GFP versus NT samples were selected. Proteins recovered in more than 200/411 IP analyses according to the contaminant repository for affinity purification (the CRAPome^39^), hence the contaminant commonly isolated or identified non-specifically by this technic, were removed from further analysis. Proteins selected according these two criteria (log2(fold change) GFP/NT>0 and not contaminant according to the CRAPome) are highlighted in green in Supp Table 2, and those showing at least 2 peptides in at least 2 replicates of the CD9-eGFP and/or the CD63-eGFP samples were used for the quantitative analysis (Supp Table 3 and Supp figure 4B). Proteins identified by at least 2 peptides identical in both CD9-eGFP and CD63-eGFP samples were considered enriched in one sample compared to the other if they showed a log2(fold change) ≥ 1 or ≤ −1 and an adjusted p-value of less than 0.05. Proteins with a log2(fold change) between −1 and 1 were considered as common to two samples. Proteins displaying peptides exclusively in CD63-eGFP or in CD9-eGFP and with at least 2 peptides in 2 replicates were listed as unique to the corresponding sample. The mass spectrometry proteomics raw data have been deposited to the ProteomeXchange Consortium via the PRIDE^40^ partner repository with the dataset identifier PXD021515 (, password: z9bnys7j).

GO term analysis was performed using the FunRich software^14,15^ using the HeLa Spatial Proteome database^13^. The hypergeometric uncorrected p-value and the percentage of proteins compared to the total number of proteins in the list were calculated by the software for each cellular component.

## Acknowledgments

We thank for fruitful discussions several team members, especially Dr Mercedes Tkach, Eleonora Grisard, Lorena Martin-Jaular and Jason Ecard, and for helpful discussions and tools, Dr P. Zimmermann, KU Leuven, Belgium and CRCM Marseille, France, Dr G. van Niel, IPN Paris, France, Dr P Benaroch, Institut Curie, Paris, France, Dr Suresh Mathivanan, LaTrobe University, Melbourne, Australia.

This work was funded by INSERM, CNRS, Institut Curie, French IdEx and LabEx (ANR-10-INSB-04, ANR-10-IDEX-0001-02 PSL, ANR-10-LABX-0038, ANR-11-LABX-0043, ANR-18-IDEX-0001 Université de Paris), grants from french ANR (ANR-18-CE13-0017-03; ANR-18-CE15-0008-01; ANR-18-CE16-0022-02), INCa (INCA-11548), Fondation ARC (PGA1 RF20180206962), FRM (FDT201904007945, EQU201903007925 and DGE20121125630), Cancéropôle Île-de-France (2013-2-EML-02-ICR-1), USA NIDA (DA040385), and a Long-Term EMBO Fellowship (ALTF 607-2015) co-funded by the European Commission FP7 (Marie Curie Actions, LTFCOFUND2013, GA-2013-609409) to J.I.V. We also acknowledge the PICT-IBiSA, member of the France-BioImaging national research infrastructure.

## Authors contribution

M. Mathieu, CT designed the study, interpreted the data, wrote the article. M Mathieu, NN, MJ, JIV, FD, D Lankar, performed experiments. M Mathieu, M Maurin, MJ analyzed the data. FV, RP, ER, GB generated plasmids or KO cells. GB, FP, designed and supervised the RUSH studies. D Loew designed and supervised the proteomic study. ER interpreted data. All authors read and corrected the article.

## Competing Interest

The authors declare no conflict of interest.

---

**Supp table 1: Quantification of the distribution of RUSH-CD63 and RUSH-CD9 constructs by immuno-electron microscopy.** RUSH-CD63-mCherry and RUSH-CD9-eGFP were labeled with antibodies against mCherry (15nm gold particles) and against GFP (10nm gold particles), at 1h or 2h after biotin addition, or at steady-state (constant presence of biotin since transfection). Ngo: observed number of gold particles. P: number of digitally generated points. Nge: expected number of gold particles. LD: labelling density (Ngo/P). RLI: relative labelling index (LD compartment/LD cell) indicates preferential labeling of the compartment when RLI >1. X2: partial χ2 = (Ngo-Nge)^2^/Nge.

**Supp table 2: Results of the quantitative mass spectrometry of the GFP samples compared to the control NT samples.** Tab1: CD63-eGFP/NT 3h, Tab2: CD9-eGFP/NT 3h, Tab3: CD63-eGFP/NT 24H, Tab4: CD9-eGFP/NT 24h. In green are shown the proteins selected as specifically immuno-isolated, by comparison to the NT samples and to the CRAPome database^39^ (see Materials and Methods for details).

**Supp table 3: Results of the quantitative mass spectrometry of the CD63-eGFP samples compared to the CD9-eGFP samples.** Tab1: All proteins used for the quantitative comparison of CD63-eGFP and CD9-eGFP EVs at 3h and 24h after biotin addition. Proteins with 1>Log2(CD9-GFP/CD63-GFP)>-1 are considered common. Tab2: Proteins enriched and unique in CD63-eGFP EVs at 3h and 24h. Log2(CD9-GFP/CD63-GFP) ≤ −1. Tab3: Proteins enriched and unique in CD9-eGFP EVs at 3h and 24h. Log2(CD9-GFP/CD63-GFP) ≥ 1. In red: membrane-associated proteins (see Materials and Methods for details of criteria used).

**Supp Figure 1:**
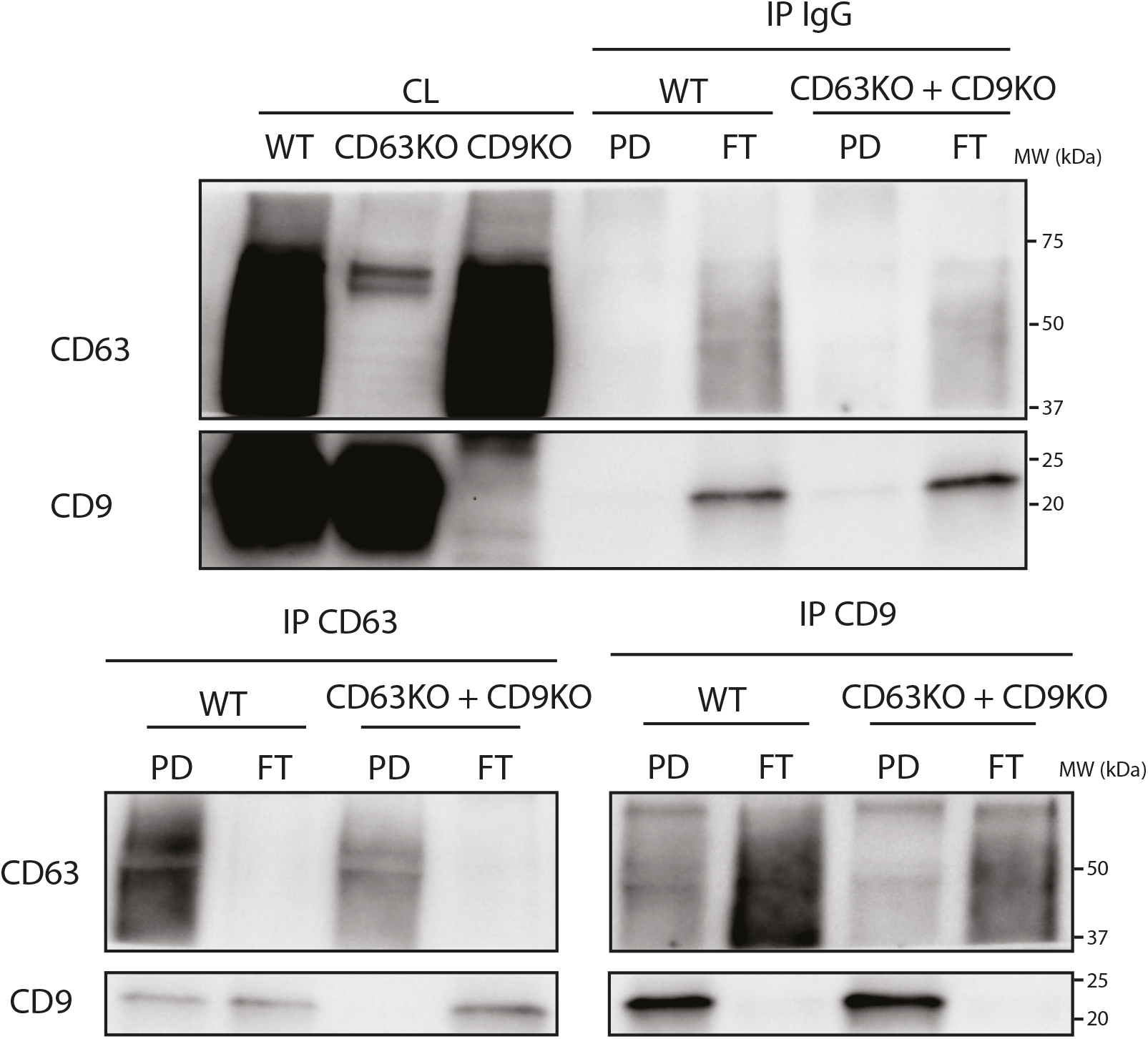
Western blot of the cell lysates (CL) of HeLa WT, CD63KO and CD9KO, and of the pull-down (PD) and flow-through (FT) after immunoprecipitation (IP) by irrelevant IgG, anti-CD63 or anti-CD9, of HeLa concentrated conditioned medium (CCM) (WT), as compared to a 1/1 mixture of HeLa CD63KO and HeLa CD9KO CCM.

**Supp figure 2:**
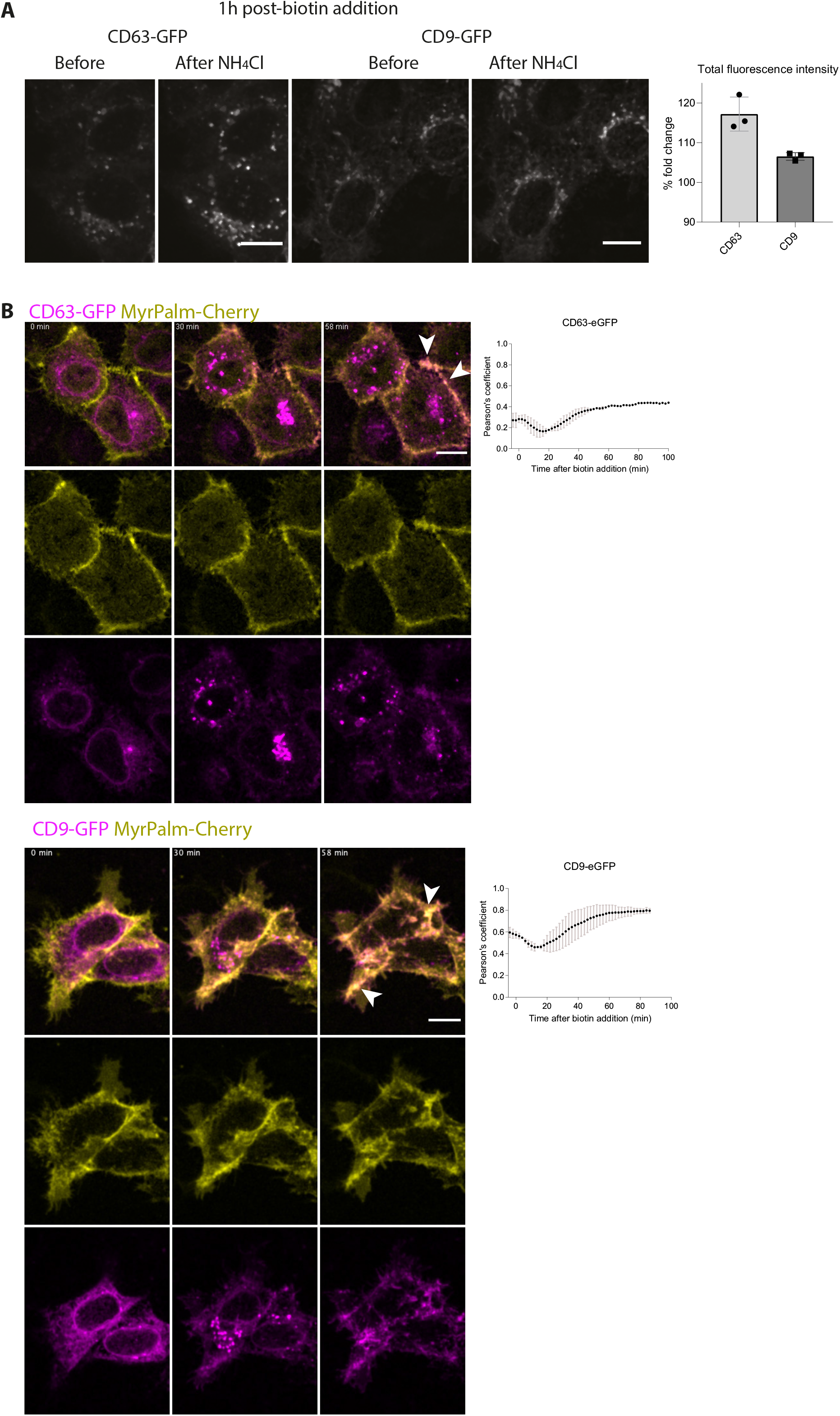
Live visualization of acidic compartments and PM by, respectively, NH_4_Cl treatment and MyrPalm-mCherry. **A)** Micrographs and quantification of live imaging of HeLa cells transfected with the RUSH plasmid CD63-eGFP before and after addition in the medium of NH_4_Cl at 50mM, 1h after biotin addition. Quantification of the fold change after NH_4_Cl addition of the total GFP fluorescence intensity. Zprojection of 11 planes. Scale bar: 5μM **B)** Micrographs of HeLa cells co-transfected with CD63 or CD9-eGFP RUSH plasmids and MyrPalm-mCherry. Biotin was added at T=0. Zprojection of 11 planes. Scale bar: 5μm. White arrows indicate some sites of co-localization. The Pearson’s co-localization coefficient between eGFP and mCherry is represented over time after biotin addition. The co-localization was quantified in one experiment for CD63 and two independent experiments for CD9.

**Supp figure 3 :**
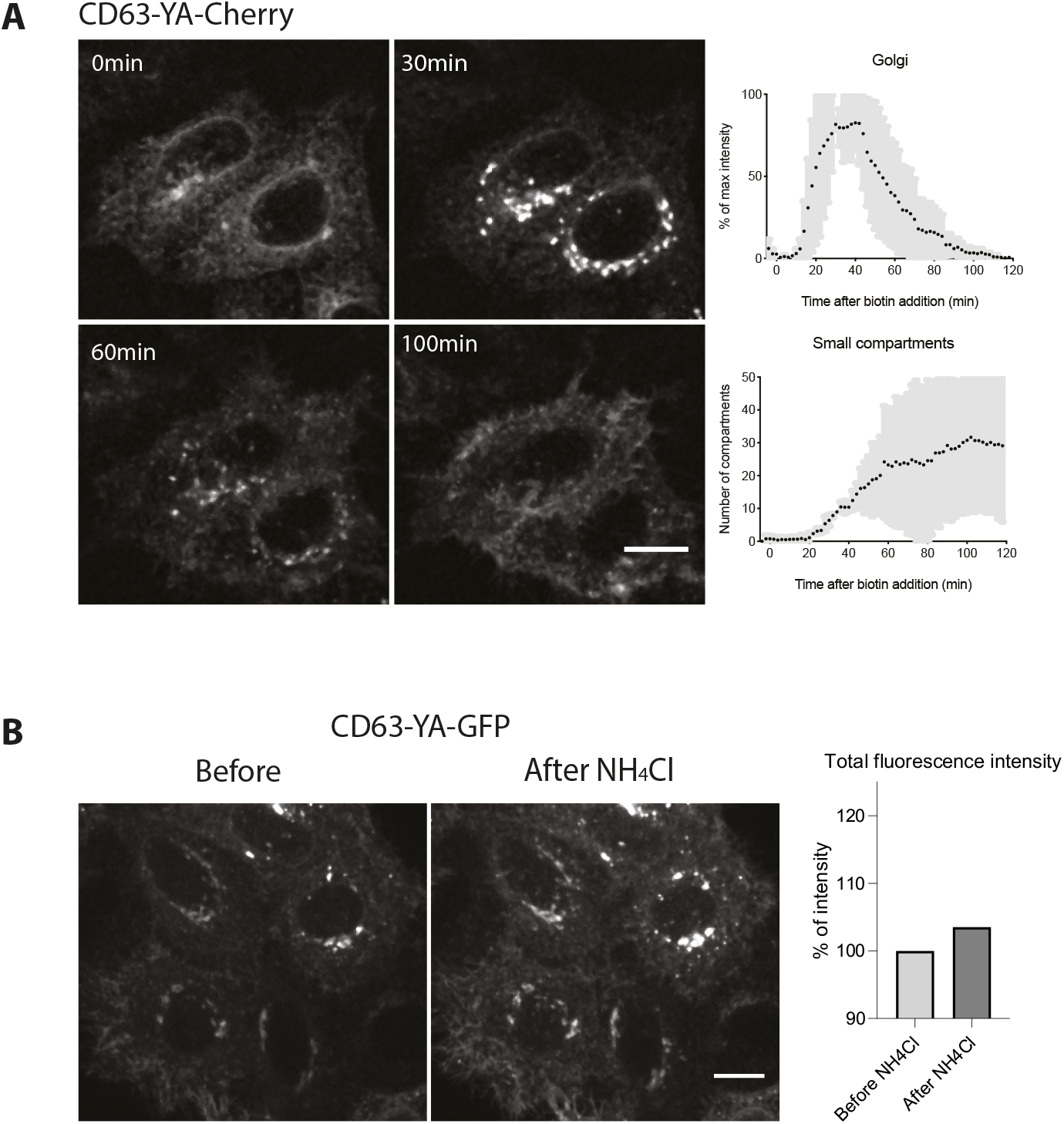
**A)** Micrographs and quantification of live imaging of HeLa cells transfected with the RUSH CD63-YA-mCherry plasmid. Biotin was added at T=0. Zprojection of 11 planes. Scale bar 5μm. Quantifications in three independent experiments. The automatically quantified Cherry fluorescence intensity in the Golgi and number of Cherry positive small compartments are represented. **B)** Extracts and quantification of live imaging of HeLa cells transfected with the RUSH plasmid CD63-YA-eGFP before and after addition in the medium of NH_4_Cl at 50mM, 1h after biotin addition. Zprojection of 11 planes. Scale bar 5μm. Quantification of the fold change after NH_4_Cl addition of the fluorescence intensity into GFP positive large compartments and of the number of GFP positive small compartments.

**Supp figure 4 :**
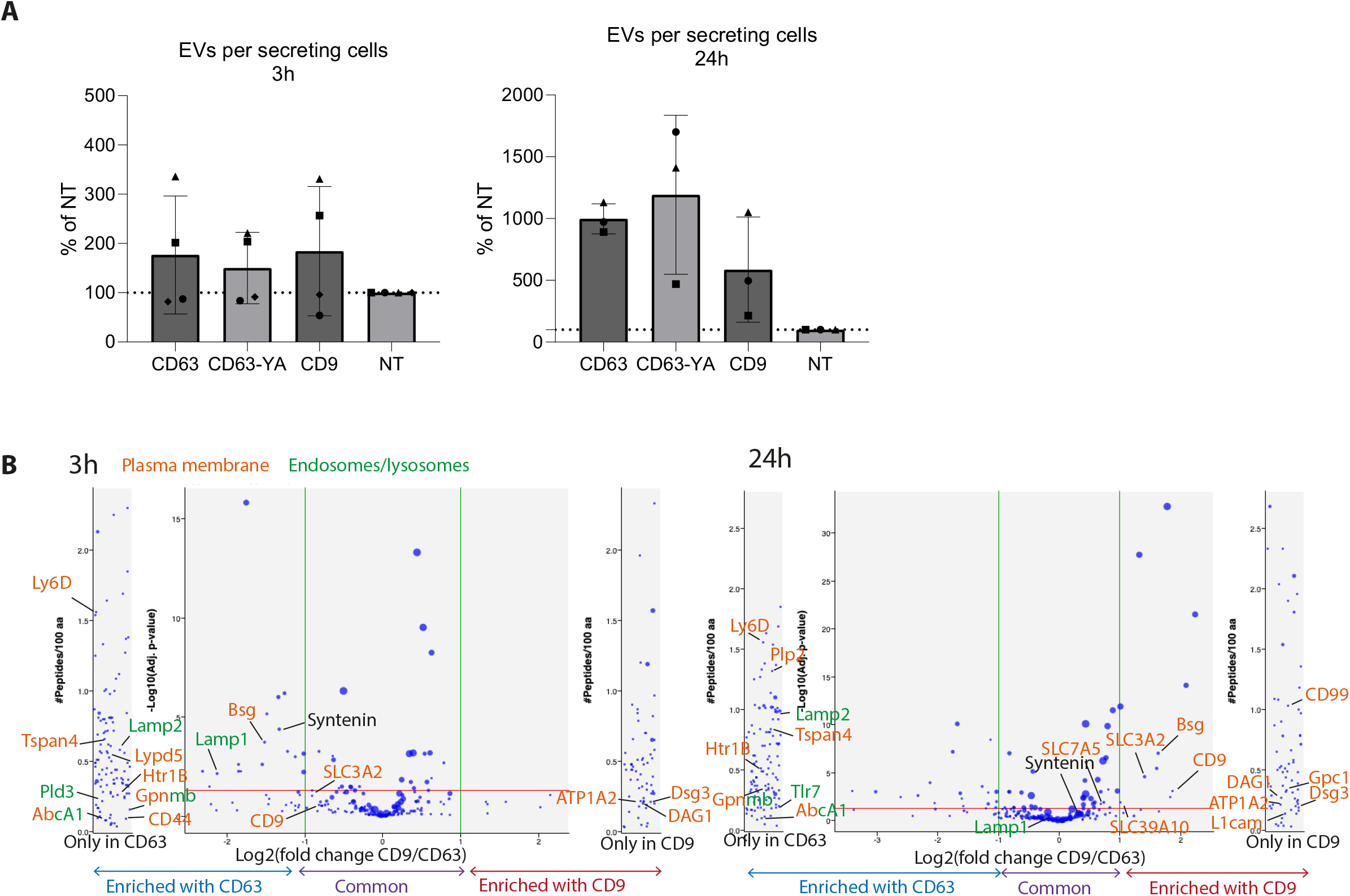
**A)** Number of EVs secreted per cell in each condition, at 3h (left) or 24h (right) after biotin addition, quantified by NTA, normalized to the NT sample. B) Volcano plots representing quantified proteins with at least 2 peptides in 2 replicates in at least one condition. Shown are the fold changes of peptide abundancy between CD63- and CD9-eGFP expressing EV samples and the p-value of this quantification, for EVs recovered 3h (left) or 24h (right) after biotin addition. Position of membrane-associated proteins and of syntenin are indicated.

**Supp figure 5 :**
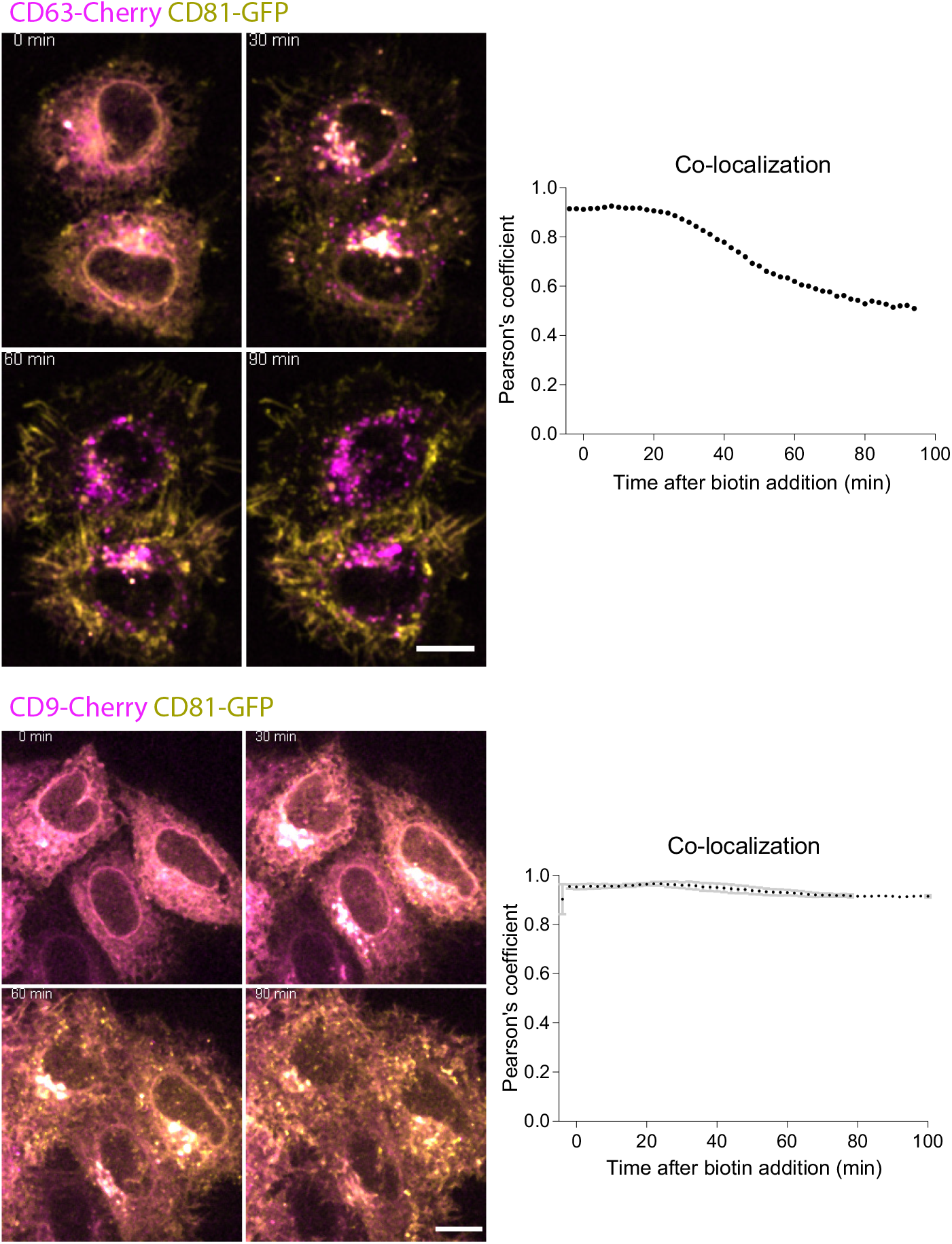
Micrographs of live imaging of HeLa cells co-transfected with the CD63-mCherry or CD9-mCherry and CD81-eGFP RUSH plasmids. Biotin was added at T=0. The Pearson’s colocalization coefficient between eGFP and mCherry is represented over time after biotin addition. Scale bar: 10μm.

## Notes

### Competing Interest Statement

The authors have declared no competing interest.

## References

1. Cocozza, F., Grisard, E., Martin-jaular, L., Mathieu, M. & Théry, C. SnapShot: Extracellular Vesicles. Cell 182, 262–262.e1 (2020).

2. Van Niel, G., D’Angelo, G. & Raposo, G. Shedding light on the cell biology of extracellular vesicles. Nat. Rev. Mol. Cell Biol. 19, 213–228 (2018).

3. Witwer, K. W. & Théry, C. Extracellular vesicles or exosomes? On primacy, precision, and popularity influencing a choice of nomenclature. J. Extracell. Vesicles 8, 1648167 (2019).

4. Mathieu, M., Martin-Jaular, L., Lavieu, G. & Théry, C. Specificities of secretion and uptake of exosomes and other extracellular vesicles for cell-to-cell communication. Nat. Cell Biol. 21, 9–17 (2019).

5. Escola, J. M. et al. Selective enrichment of tetraspan proteins on the internal vesicles of multivesicular endosomes and on exosomes secreted by human B-lymphocytes. J Biol Chem 273, 20121–20127 (1998).

6. Thery, C. et al. Molecular characterization of dendritic cell-derived exosomes. Selective accumulation of the heat shock protein hsc73. J Cell Biol 147, 599–610 (1999).

7. Kowal, J. et al. Proteomic comparison defines novel markers to characterize heterogeneous populations of extracellular vesicle subtypes. Proc. Natl. Acad. Sci. 113, E968–E977 (2016).

8. Boncompain, G. et al. Synchronization of secretory protein traffic in populations of cells. Nat. Methods 9, 493–498 (2012).

9. Verweij, F. J. et al. Quantifying exosome secretion from single cells reveals a modulatory role for GPCR signaling. J. Cell Biol. 217, 1129–1142 (2018).

10. Rous, B. A. et al. Role of Adaptor Complex AP-3 in Targeting Wild-Type and Mutated CD63 to Lysosomes. Mol. Biol. Cell 13, 1071–1082 (2002).

11. Edgar, J. R., Manna, P. T., Nishimura, S., Banting, G. & Robinson, M. S. Tetherin is an exosomal tether. Elife 5, (2016).

12. Cashikar, A. G. & Hanson, P. I. A cell-based assay for CD63-containing extracellular vesicles. PLoS One 14, e0220007 (2019).

13. Itzhak, D. N., Tyanova, S., Cox, J. & Borner, G. H. H. Global, quantitative and dynamic mapping of protein subcellular localization. Elife 5, e16950 (2016).

14. Pathan, M. et al. FunRich: An open access standalone functional enrichment and interaction network analysis tool. Proteomics 15, 2597–2601 (2015).

15. Pathan, M. et al. A novel community driven software for functional enrichment analysis of extracellular vesicles data. J. Extracell. Vesicles 6, 1321455 (2017).

16. Baietti, M. F. et al. Syndecan–syntenin–ALIX regulates the biogenesis of exosomes. Nat. Cell Biol. 14, 677–685 (2012).

17. Trajkovic, K. et al. Ceramide triggers budding of exosome vesicles into multivesicular endosomes. Science (80-.). 319, 1244–1247 (2008).

18. Menck, K. et al. Microvesicles mediate Breast cancer invasion through Glycosylated EMMPRIN. J. Mol. Cell Biol. 7, 143–153 (2015).

19. Minciacchi, V. R. et al. Large oncosomes contain distinct protein cargo and represent a separate functional class of tumor-derived extracellular vesicles. Oncotarget 6, 11327–11341 (2015).

20. Keerthikumar, S. et al. Proteogenomic analysis reveals exosomes are more oncogenic than ectosomes. Oncotarget 6, 15375–15396 (2015).

21. Zhang, H. et al. Identification of distinct nanoparticles and subsets of extracellular vesicles by asymmetric flow field-flow fractionation. Nat. Cell Biol. 20, 332–343 (2018).

22. Jeppesen, D. K. et al. Reassessment of Exosome Composition. Cell 177, 428–445.e18 (2019).

23. Mannioni, B. A. et al. The light chain of CD98 is identified as E16/TA1 protein. J. Biol. Chem. 273, 33127–33129 (1998).

24. Xu, D. & Hemler, M. E. Metabolic activation-related CD147-CD98 complex. Mol. Cell. Proteomics 4, 1061–1071 (2005).

25. Ip, H. & Sethi, T. CD98 signals controlling tumorigenesis. Int. J. Biochem. Cell Biol. 81, 148–150 (2016).

26. Muramatsu, T. Basigin (CD147), a multifunctional transmembrane glycoprotein with various binding partners. J. Biochem. 159, 481–490 (2016).

27. Cantor, J. M. & Ginsberg, M. H. CD98 at the crossroads of adaptive immunity and cancer. J. Cell Sci. 125, 1373–1382 (2012).

28. Arendt, B. K., Walters, D. K., Wu, X., Tschumper, R. C. & Jelinek, D. F. Multiple myeloma dell-derived microvesicles are enriched in CD147 expression and enhance tumor cell proliferation. Oncotarget 5, 5686–5699 (2014).

29. Abache, T. et al. The transferrin receptor and the tetraspanin web molecules CD9, CD81, and CD9P-1 are differentially sorted into exosomes after TPA treatment of K562 cells. J. Cell. Biochem. 102, 650–664 (2007).

30. Palokangas, H., Ying, M., Väänänen, K. & Saraste, J. Retrograde transport from the pre-Golgi intermediate compartment and the Golgi complex is affected by the vacuolar H+-ATPase inhibitor bafilomycin A1. Mol. Biol. Cell 9, 3561–3578 (1998).

31. Ran, F. A. et al. Genome engineering using the CRISPR-Cas9 system. Nat. Protoc. 8, 2281–2308 (2013).

32. Charrin, S. et al. Rapid Isolation of Rare Isotype-Switched Hybridoma Variants: Application to the Generation of IgG2a and Exosome Marker. Antibodies 9, E29 (2020).

33. Le Naour, F. et al. Profiling of the tetraspanin web of human colon cancer cells. Mol. Cell. Proteomics 5, 845–857 (2006).

34. Slot, J. W. & Geuze, H. J. Cryosectioning and immunolabeling. Nat. Protoc. 2, 2480–2491 (2007).

35. Mayhew, T. M. Mapping the distributions and quantifying the labelling intensities of cell compartments by immunoelectron microscopy: Progress towards a coherent set of methods. J. Anat. 219, 647–660 (2011).

36. Poullet, P., Carpentier, S. & Barillot, E. myProMS, a web server for management and validation of mass spectrometry-based proteomic data. Proteomics 7, 2553–2556 (2007).

37. The, M., MacCoss, M. J., Noble, W. S. & Käll, L. Fast and Accurate Protein False Discovery Rates on Large-Scale Proteomics Data Sets with Percolator 3.0. J. Am. Soc. Mass Spectrom. 27, 1719–1727 (2016).

38. Valot, B., Langella, O., Nano, E. & Zivy, M. MassChroQ: a versatile tool for mass spectrometry quantification. Proteomics 11, 3572–3577 (2011).

39. Mellacheruvu, D. et al. The CRAPome: a contaminant repository for affinity purification-mass spectrometry data. Nat Methods 10, 730–736 (2013).

40. Perez-Riverol, Y. et al. The PRIDE database and related tools and resources in 2019: Improving support for quantification data. Nucleic Acids Res. 47, D442–D450 (2019).

